# Cellular resolution anatomical and molecular atlases for prenatal human brains

**DOI:** 10.1101/2021.07.14.452297

**Authors:** Song-Lin Ding, Joshua J. Royall, Phil Lesnar, Benjamin A.C. Facer, Kimberly A. Smith, Yina Wei, Kristina Brouner, Rachel A. Dalley, Nick Dee, Tim A. Dolbeare, Amanda Ebbert, Ian A. Glass, Katie Glattfelter, Nika H. Keller, Felix Lee, Tracy A. Lemon, Julie Nyhus, Julie Pendergraft, Robert Reid, Melaine Sarreal, Nadiya V. Shapovalova, Aaron Szafer, John W. Phillips, Susan M. Sunkin, John G. Hohmann, Allan R. Jones, Michael J. Hawrylycz, Patrick R. Hof, Lydia Ng, Amy Bernard, Ed S. Lein

## Abstract

Increasing interest in studies of prenatal human brain development, particularly using new single-cell genomics and anatomical technologies to create cell atlases, creates a strong need for accurate and detailed anatomical reference atlases. In this study, we present two cellular-resolution digital anatomical atlases for prenatal human brain at post-conceptional weeks (PCW) 15 and 21. Both atlases were annotated on sequential Nissl-stained sections covering brain-wide structures on the basis of combined analysis of cytoarchitecture, acetylcholinesterase staining and an extensive marker gene expression dataset. This high information content dataset allowed reliable and accurate demarcation of developing cortical and subcortical structures and their subdivisions. Furthermore, using the anatomical atlases as a guide, spatial expression of 37 and 5 genes from the brains respectively at PCW 15 and 21 was annotated, illustrating reliable marker genes for many developing brain structures. Finally, the present study uncovered several novel developmental features, such as the lack of an outer subventricular zone in the hippocampal formation and entorhinal cortex, and the apparent extension of both cortical (excitatory) and subcortical (inhibitory) progenitors into the prenatal olfactory bulb. These comprehensive atlases provide useful tools for visualization, targeting, imaging and interpretation of brain structures of prenatal human brain, and for guiding and interpreting the next generation of cell census and connectome studies.

## 1. INTRODUCTION

Anatomical brain atlases are essential tools for visualizing, integrating and interpreting experimental data about brain structure, function, circuits, cell types and structure-function-behavior relationships (Evans et al., 2012; Wang et al., 2020). We previously generated brain-wide detailed microarray-based transcriptomic atlases for the prenatal human brain at post-conceptional weeks (PCW) 15, 16 and 21 (Miller et al., 2014), and single cell genomic studies are now increasingly profiling prenatal brains to define cellular diversity, developmental trajectories and gene regulatory mechanisms (Nowakowski et al., 2017; Fan et al., 2020; Eze et al., 2021). To provide an anatomical and ontological framework for these prior and future studies of human brain development, here we aimed to create detailed and accurate reference atlases that densely sample the whole developing brain at PCW 15 and 21. These prenatal human brain atlases can also be important tools to guide increasing neuroimaging studies of prenatal human brains and developmental deficits and malformations (Oishi et al., 2018; Kostović et al., 2019).

Two highly detailed comprehensive anatomical atlases are available for the adult human brain (Ding et al. 2016; Mai et al., 2016). Fewer anatomical references are available for developing human brains, and especially for prenatal stages. Only one series of prenatal human brain atlases is available, generated on a limited set of Nissl-stained sections from different brain specimens (Bayer & Altman, 2003; 2005; 2006). While heroic efforts at the time, these atlases have relatively low sampling density, are limited to Nissl or hematoxylin-eosin stains, and have fewer structural annotations than the adult atlases. For example, only 15 and 13 coronal sections were annotated for the human brain atlases from prenatal weeks (PW) 13.5 and 17, respectively (Bayer & Altman, 2005). Furthermore, certain developmental stages, such as PW 15 and 16, are not available in this atlas series.

In this study we aimed to create a plate-based atlas with coverage of essentially all anatomical structures, a complete developmental structural ontology, and a high information content gene expression analysis that allows accurate structural delineation. Whole brain serial sectioning was performed on each brain, with interdigitated histochemistry and *in situ* hybridization (ISH) spanning the entire brain specimens, which were scanned at 1µm/pixel resolution. We annotated representative Nissl-stained coronal sections spanning the brain (46 sections for the PCW 15 brain and 81 sections for the PCW 21 brain) based on a combined analysis of cytoarchitecture, acetylcholinesterase (AChE) staining and expression patterns of selected genes from the same brain. Annotations were also performed on a series of the *in situ* hybridization (ISH) images that often delineate particular structures well (sections for 37 and 5 selected genes from the PCW 15 and 21 brains, respectively). Finally, these atlases are presented as freely accessible online interactive data resources (www.brain-map.org or www.brainspan.org).

## 2. MATERIALS AND METHODS

### 2.1 Prenatal human brain specimens

Two post-mortem human brain specimens at PCW 15 (male; Caucasian) and PCW 21 (female; Asian), respectively, were used for generation of anatomical and molecular atlases. Both specimens were procured from Laboratory of Developmental Biology at the University of Washington, Seattle, USA. All work was performed according to guidelines for the research use of human brain tissue and with approval by the Human Investigation Committees and Institutional Ethics Committees of University of Washington. Appropriate written informed consent was obtained and all available non-identifying information was recorded for each specimen. Both brains showed normal appearance and high RNA quality with an average RNA integrity number of 8 and 9 for PCW 15 and 21, respectively. Both brains were bisected, and the left hemisphere was used for DNA microarray analysis (see Miller et al, 2014) and right hemisphere was used for histology and ISH stains. For the right hemisphere, two and four coronal slabs were cut for the PCW 15 and 21 brains, respectively, based on the size of the hemisphere. These slabs were frozen in isopentane chilled to -50°C and stored at -80°C until sectioning. Serial sectioning was performed through the whole hemisphere, slab by slab. Nissl, acetylcholinesterase (AChE) and ISH stains for 43 gene probes were carried out on sequential series of sections (see below). For both stages, sequential sections for Nissl and AChE stain were regularly spaced and flanked by series of ISH for marker genes. All histology and ISH sections were digitally scanned at 1.0 µm/pixel. To generate anatomical atlases for PCW 15 and 21 brains, 46 out of the 115, and 81 out of the 174 Nissl-stained sequential sections were selected, respectively.

### 2.2 Nissl staining

After sectioning 20 μm-thick sections in the coronal plane from an entire hemisphere of the specimens, slides were baked at 37°C for 1 to 5 days and were removed 5 to 15 minutes prior to staining. Sections were defatted with xylene or the xylene substitute Formula 83, and hydrated through a graded series containing 100, 95, 70, and 50% ethanol. After incubation in water, the sections were stained in 0.213% thionin, then differentiated and dehydrated in water and a graded series containing 50, 70, 95, and 100% ethanol. Finally, the slides were incubated in xylene or xylene substitute Formula 83, and coverslipped with the mounting agent DPX. After drying, the slides were analyzed microscopically to ensure staining quality.

### 2.3 AChE staining

A modified AChE protocol was used to help delineate subcortical structures at high resolution. AChE staining was performed using a direct coloring thiocholine method combined with a methyl green nuclear counterstain to improve tissue visibility (Karnovsky & Roots, 1964). Glass slides with fresh-frozen tissue sections were removed from 4°C, allowed to equilibrate to room temperature, fixed in 10% neutral buffered formalin and washed briefly in ultra-pure water. Sections were then incubated for 30 minutes in a solution of acetylthiocholine iodide, sodium citrate, cupric sulfate, and potassium ferricyanide in a 0.1M sodium acetate buffer (pH 6.0), washed in 0.1M Tris-HCl buffer (pH 7.2), incubated with 0.5% diaminobenzidine in 0.1M Tris-HCl with 0.03% hydrogen peroxide. Slides were incubated in 0.2% methyl green, briefly dipped in 100% ethanol, cleared with Formula 83 and coverslipped with DPX.

### 2.4 ISH staining

A colorimetric, digoxigenin-based method for labeling target mRNA was used to detect gene expression on human prenatal tissue sections with 43 selected genes (see Lein et al., 2007). These genes include canonical morphological and cell type markers and disease-related genes associated with neocortical development. Gene selection was preferential towards data available through the Allen Developing Mouse Brain Atlas (Thompson et al., 2014), allowing a direct phylogenetic comparison of gene expression patterns between mouse and human. Gene lists and details of the ISH process are available online (http://help.brain-map.org/display/devhumanbrain/Documentation). Gene list is also shown in the legend of appendix 2.

### 2.5 Digital imaging and image processing

Digital imaging of the stained slides was done using a ScanScope XT (Aperio Technologies Inc., Vista, CA) with slide autoloader. The final resolution of the images was 1 μm/pixel. All images were databased and preprocessed, then subjected to quality control (QC) to ensure optimal focus and that no process artifacts were present on the slide images. Images that passed this initial QC were further assessed to ensure that the staining data were as expected. Once all QC criteria were met, images became available for annotation of anatomical structures.

### 2.6 Generation of whole-brain structure ontology

To generate a unifying hierarchical ontology for both developing and adult human brains with each structure having a unique identification code, we first subdivided the brain into three major parts: forebrain, midbrain and hindbrain. Under each major part, we created four main branches: gray matter, white matter, ventricles and surface structures (e.g., cortical sulci and gyri). Under the gray matter branches two types of brain structures were separated: transient and permanent ones. Under the transient structures we listed all structures that only appear during development and not exist in adult brain (see Table 1 for detailed transient structures). Under the permanent structures we listed all structures that exist in both developing and adult brains (for details see Table 3 of Ding et al., 2016). Table 1 also lists abbreviations for the main brain structures shown in this study.

**Table 1.** Abbreviations and ontology of brain structures (not attached due to large file size)

### 2.7 Creation of prenatal human brain atlases

For the specimen at PCW 15, a total of 115 Nissl-stained sections were produced at 1.04 mm spacing. For annotation 46 slides were chosen, including 23 from slab 1 (∼1 mm sampling density for the first 7 Nissl-stained levels, ∼0.5 mm for the remaining 16 ones), and 22 from slab two (∼0.5 mm sampling density for the first 16 Nissl-stained levels, ∼1 mm for the remaining 6 ones), and a single additional section effectively between slabs one and two. For the specimen at PCW 21, four slabs were generated due to its larger size than the PCW15 brain. Each of these four slabs were sectioned into 174 Nissl-stained sections with 3 per 1.2 mm. A total of 81 - stained levels were chosen for annotation for anatomical atlas of this stage including 13 from slab 1 (∼1.2 mm sampling density), 32 from slab 2 (∼0.5 mm sampling density), 22 from slab 3 (∼0.5 mm sampling density for the first 16, ∼1.2 mm sampling density for the remaining 6), and 14 from slab 4 (∼1.2 mm sampling density). In general, the slabs containing smaller brain structures (usually subcortical regions) were sampled more densely to avoid missing small structures. The position of each section in a given slab was marked.

Annotation of the present brain atlases was performed similarly to that of our digital adult human brain atlas (Ding et al., 2016). Briefly, annotation drawings were done on printouts of the Nissl-stained sections and then digitally scanned. Digital cartographic translation of expert-delineated Nissl printouts was performed using Adobe Creative Suite 5. The resulting vector graphics were then converted to Scalable Vector Graphics (SVG). Each polygon was then associated with a structure from the ontology (see **Table 1**). Collating polygons in this way allows the flexibility to create various presentation modes (e.g., with or without colorization and transparency). The brain structures were colorized to assist users with identifying structures across different sections (see **Appendices 1 and 3**). Gross ontological groups (“parents”) were assigned hues from a range of the color spectrum. Each structure within a given parent group (“child”) was given a variation of the parent hue according to its relative cellular contrast in Nissl stain. The following general principle was applied: the higher the density, the deeper the shade (i.e., addition of black to hue); the lower the density, the deeper the tint (i.e., addition of white to hue). Large parent groups (e.g., thalamus) were assigned uniformly light variations of their principal hues to provide a visually subtle, cohesive backdrop for component substructures, which often exhibit a range of relative cellular contrasts (reflected by shades and tints). To create gene expression atlases for PCW 15 and 21 brains, we applied annotations from the anatomical atlases for each age onto the interleaved coronal ISH sections for 37 (PCW15) and 5 (PCW21) genes out of 43 (see **Appendices 2 and 4)**. The workflow for generation of the prenatal human brain atlases is similar to the one described in our adult human brain atlas (Ding et al., 2016) and is briefly summarized in **Figure 1**.

**Fig. 1.**
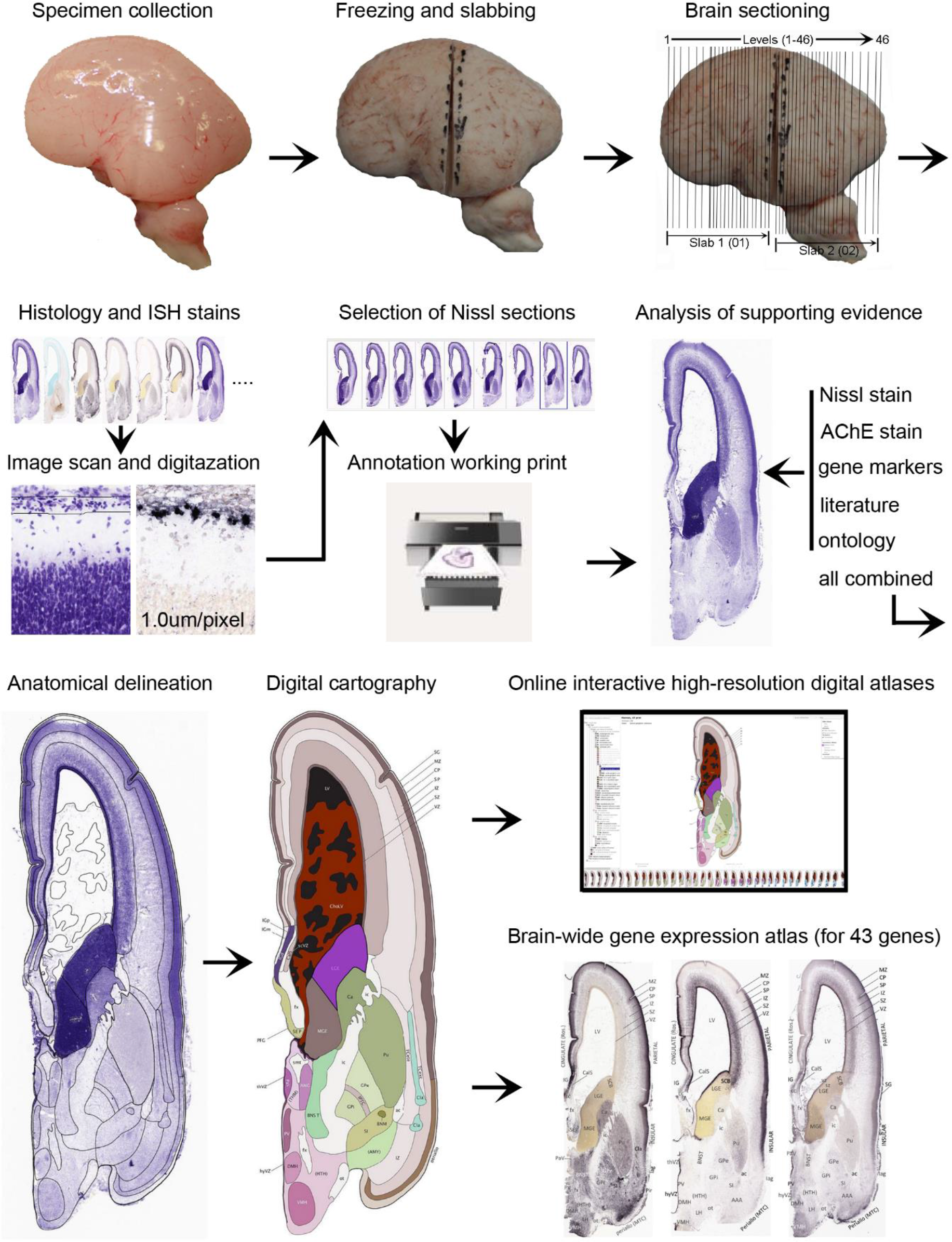
Workflow for atlas generation.

## 3. RESULTS

### 3.1 Structural annotation of histological and molecular prenatal human brain datasets

To generate accurate and detailed anatomical brain atlases, we performed both histological stains (Nissl and AChE) and ISH for 43 gene probes on sequential sets of coronal cryosections from right hemisphere of two mid-gestation brains (PCW 15 and 21). With the anatomical atlases as a guide, we also annotated the spatial expression of 37 and 5 genes in the brain at PCW 15 and 21, respectively; these are treated as prenatal molecular brain atlases. The anatomical and molecular atlases for the brain at PCW 15 are presented in **Appendices 1** and **2**, respectively. The similarly generated anatomical and molecular atlases for the brain at PCW 21 are presented in **Appendices 3** and **4**. All appendices have online links for cellular resolution histology and ISH images (1.0 µm/pixel). Example plates of annotated anatomical atlases from the two brains are shown in **Figure 2** (where a and b designate PCW 15 and PCW 21, respectively). Delineation of anatomical boundaries of different cortical layers and brain regions are detailed below with emphasis mainly on the brain at PCW 15 although some major molecular features from the brain at PCW 21 are also described for comparison.

**Fig. 2.**
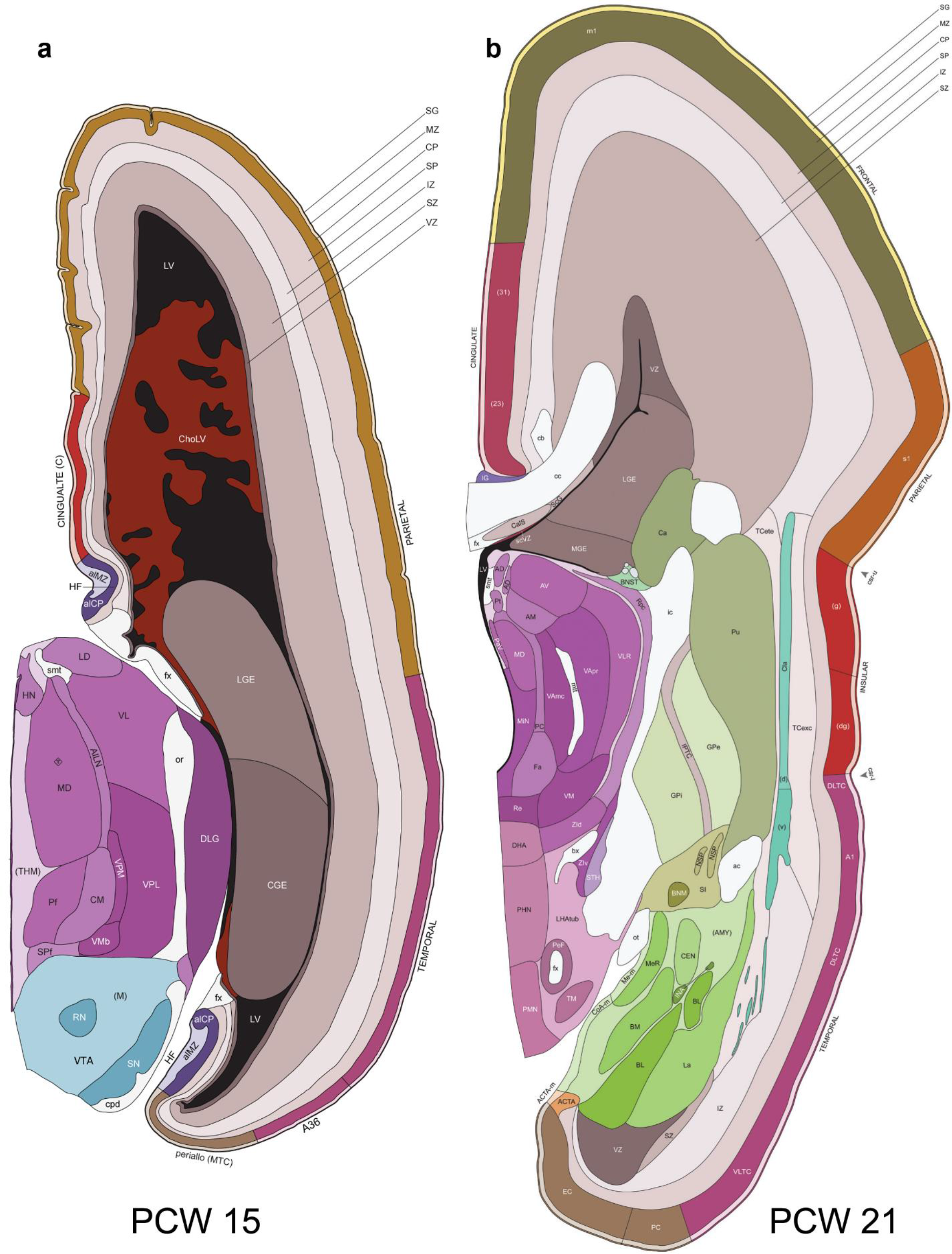
Example of anatomical atlas plates from PCW 15 (a) and 21 (b) brains.

### 3.2 Delineation of prenatal neocortical layers

#### PCW 15

In Nissl preparations, seven neocortical layers can be generally identified. For example, from the pia to the lateral ventricle (LV) of the medial occipital cortex, these layers include the subpial granular zone (SG), marginal zone (MZ), cortical plate (CP), subplate (SP), intermediate zone (IZ), subventricular zone (SZ) and ventricular zone (VZ) (**Fig. 3a**). The SZ can be further subdivided into less densely packed outer and more densely packed inner parts (SZo and SZi, respectively) with SZi adjoining VZ, which is the most densely packed zone near the LV (**Fig. 3a**). At PCW15, the outer fiber zone (OF) begins to appear in the outermost part of the SZo, deep to the IZ. To confirm and accurately to delineate the developing neocortical layers, we analyzed the large set of ISH data described above and found that many genes display layer-specific expression patterns. For instance, in the medial occipital/visual cortex (**Fig. 3**), *PAX6* and *TBR2* (*EOMES*) are selectively expressed in the proliferative zones VZ and SZ with strongest *PAX6* and *TBR2* expression in VZ and deep SZ (SZd, near VZ), respectively (**Fig. 3b, c**). SZd was sometimes termed as the border zone (BZ) between SZ and VZ. Interestingly, inner part of the VZ (VZi, near LV) does not show *TBR2* expression (**Fig. 3c**). *VIM*, *SOX2* and *FABP7* are also dominantly expressed in VZ and SZ (VZ>SZ) but with weak expression in CP (**Fig. 3d, g, h**). In contrast, *GRIK2* and *SATB2* are selectively expressed in the postmitotic zones IZ/OF, SP and CP. Specifically, *GRIK2* is dominantly expressed in SP and deep CP with weak expression in IZ while *SATB2* is mainly expressed in IZ/OF with weak expression in SP and CP (**Fig. 3e, f**). Some genes (e.g. *ENC1*) are strongly expressed in both proliferative (VZ, SZ) and post-mitotic (CP) zones (**Fig. 3i**).

**Fig. 3.**
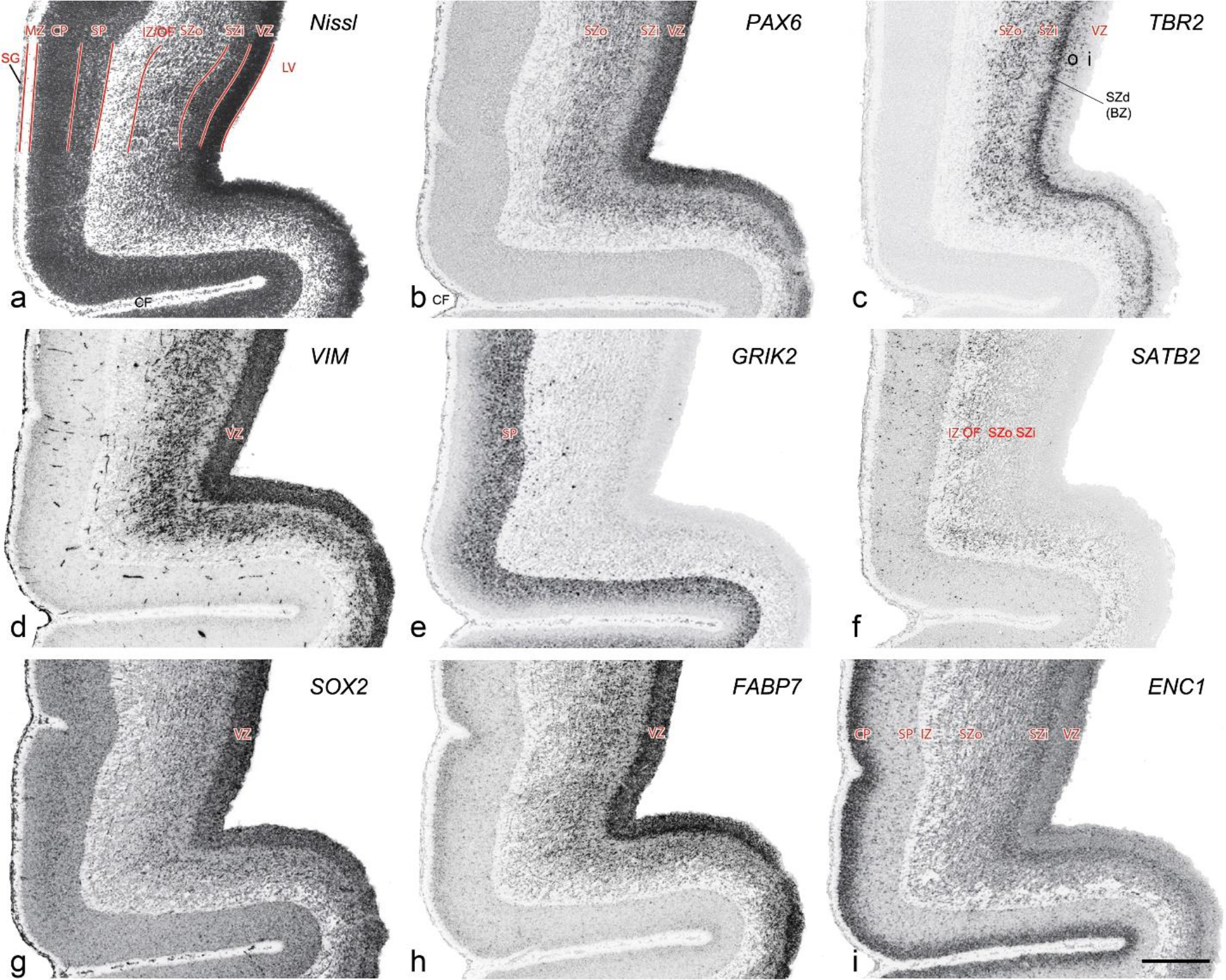
Molecular marker expression in medial occipital cortex at PCW15. (a). A Nissl-stained section showing the lamination of the cortex near the calcarine figure (CF). (b-i). Expression patterns of *PAX6* (b), *TBR2* (c), *VIM* (d), *GRIK2* (e), *SATB2* (f), *SOX2* (g), *FABP7* (h) and *ENC1* (i). Note that a dense zone of *TBR2* expression at the border between SZi and VZ is termed deep SZ zone (SZd) or border zone (BZ). The thickness ratio of SZo to VZ is about 3:1. SG, subpial granular zone; MZ, marginal zone; CP, cortical plate; SP, subplate; IZ, intermediate zone; OF, outer fiber zone; SZ, subventricular zone; SZo and SZi, outer and inner SZ; VZ, ventricular zone; LV, lateral ventricle. These terms apply to main text and all related figures below. Scale bar: 400 µm in (i) for all panels.

To examine whether more anterior neocortical regions display similar or different laminar organization we investigated the same set of genes expressed in the dorsomedial frontal neocortex (**Fig. 4**). In this region, all the cortical layers are visible with relatively thick SP compared to CP (**Fig. 4a**). Compared to the occipital region, the existence of the OF in the frontal cortex is more visible and thicker. The OF contains many radially oriented SZ cells (***Fig. 4a***) and is thus included in the outermost part of SZo in the present atlas (i.e., the OF is not annotated separately from the SZo). In the frontal cortex, *VIM*, *SOX2*, *PAX6* and *TBR2* have similar expression patterns as in the occipital cortex (e.g. **Fig. 4a-c**). Compared to the occipital cortex, *FABP7* shows stronger expression in CP (**Fig. 4d**) although expression in other zones is comparable to the occipital cortex. *LMO4* is selectively expressed in the CP (**Fig. 4e**) while this expression in the occipital cortex is not obvious (see **Appendix 2**). *GRIK2* display strong expression in CP, SP and IZ of frontal cortex (**Fig. 4g**), while it is mainly expressed in SP of occipital cortex (**Fig. 3e**). *SATB2* in the frontal cortex has strong expression in CP, SP, IZ and OF (**Fig. 4h**), while only the OF has strong expression in the occipital cortex. In addition, *ETV1* expression appears in the deep CP (**Fig. 4f**), but is not detected in the occipital cortex (not shown). The differential gene expression could reflect differential maturation across the cortex as well as regional differences. Additional data on later stages are needed to address this issue.

**Fig. 4.**
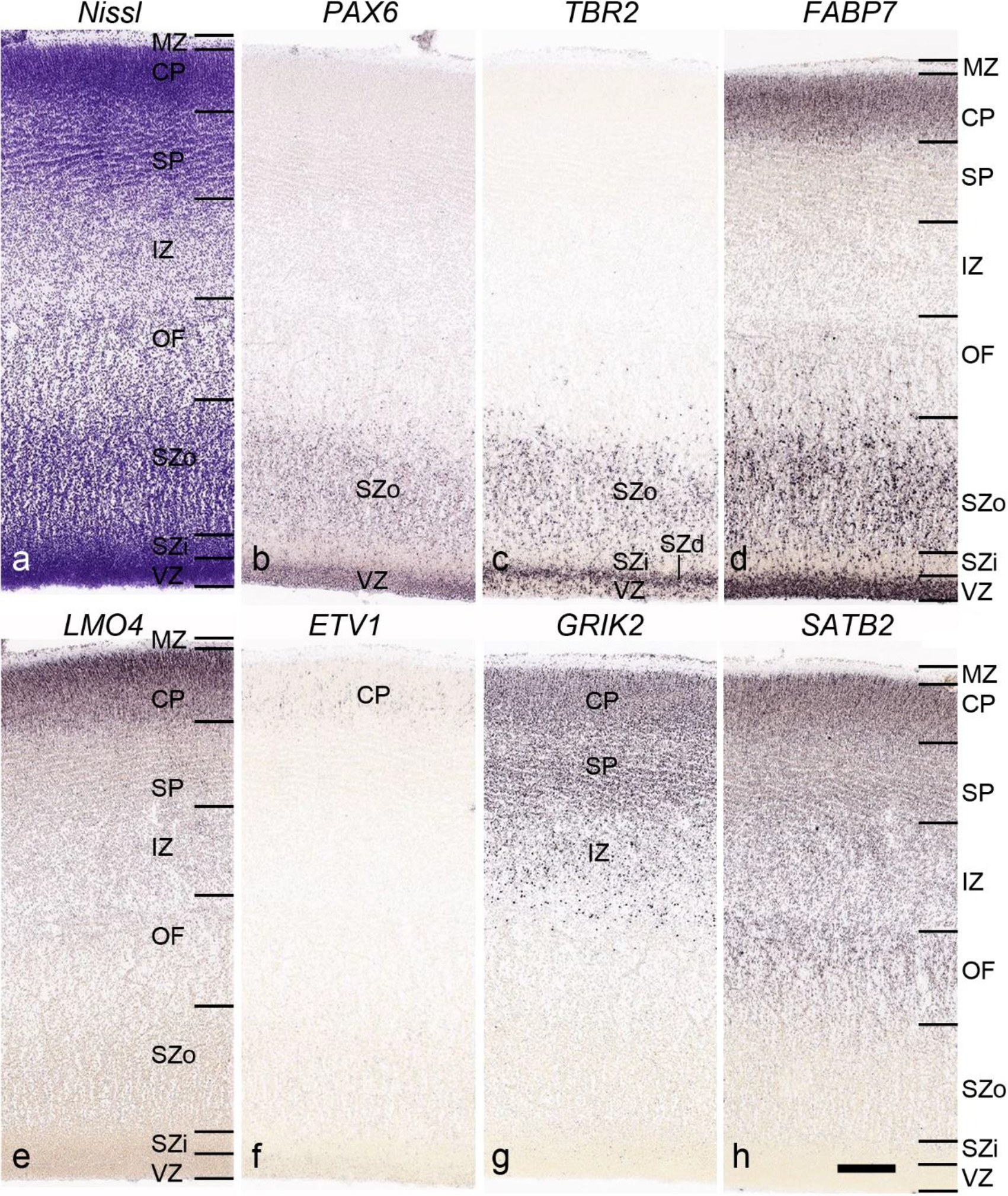
Gene expression in medial frontal cortex at PCW 15. (a). Nissl-stained section showing the laminar organization of the cortex. (b-h). Expression patterns of *PAX6* (b), *TBR2* (c), *FABP7* (d), *LMO4* (e), *ETV1* (f), *GRIK2* (g) and *SATB2* (h). Note that the thickness ratio of SZo to VZ is about 5:1. Scale bar: 330 µm in (h) for all panels.

#### PCW 21

On Nissl-stained sections, all the cortical layers that appeared at PCW 15 can be identified at PCW21, although changes in their relative thickness are observed. In the occipital/visual cortex, the thickness of CP and SZo is greatly increased and outer and inner CP (CPo and CPi) are distinguishable (**Fig. 5a**). A major feature of the neocortex at PCW 21 is the clear presence of the inner fiber zone (IF), which is a cell-sparse zone located between SZo and SZi (**Fig. 5a**). Furthermore, the boundaries of the primary visual cortex (V1) are obvious based on gene expression boundaries, but not yet discernable based on Nissl staining (**Fig. 5b-d**). *NPY* expression is mainly located in the SP of all neocortical regions and in the middle layers (future layers 3 and 4) of V1 (**Fig. 5e**). Strong *Pax6*, *TRB2* and *VIM* expression is restricted in the proliferative zones (SZo, SZi and VZ), similar to the findings from PCW 15. The OF and IF at PCW 21 still contain a lot of SZo cells and thus are included in the SZ (SZo) in our atlas plates (**Appendix 3**). *SST* is an additional marker for the SP at PCW21 (**Appendix 4**).

**Fig. 5.**
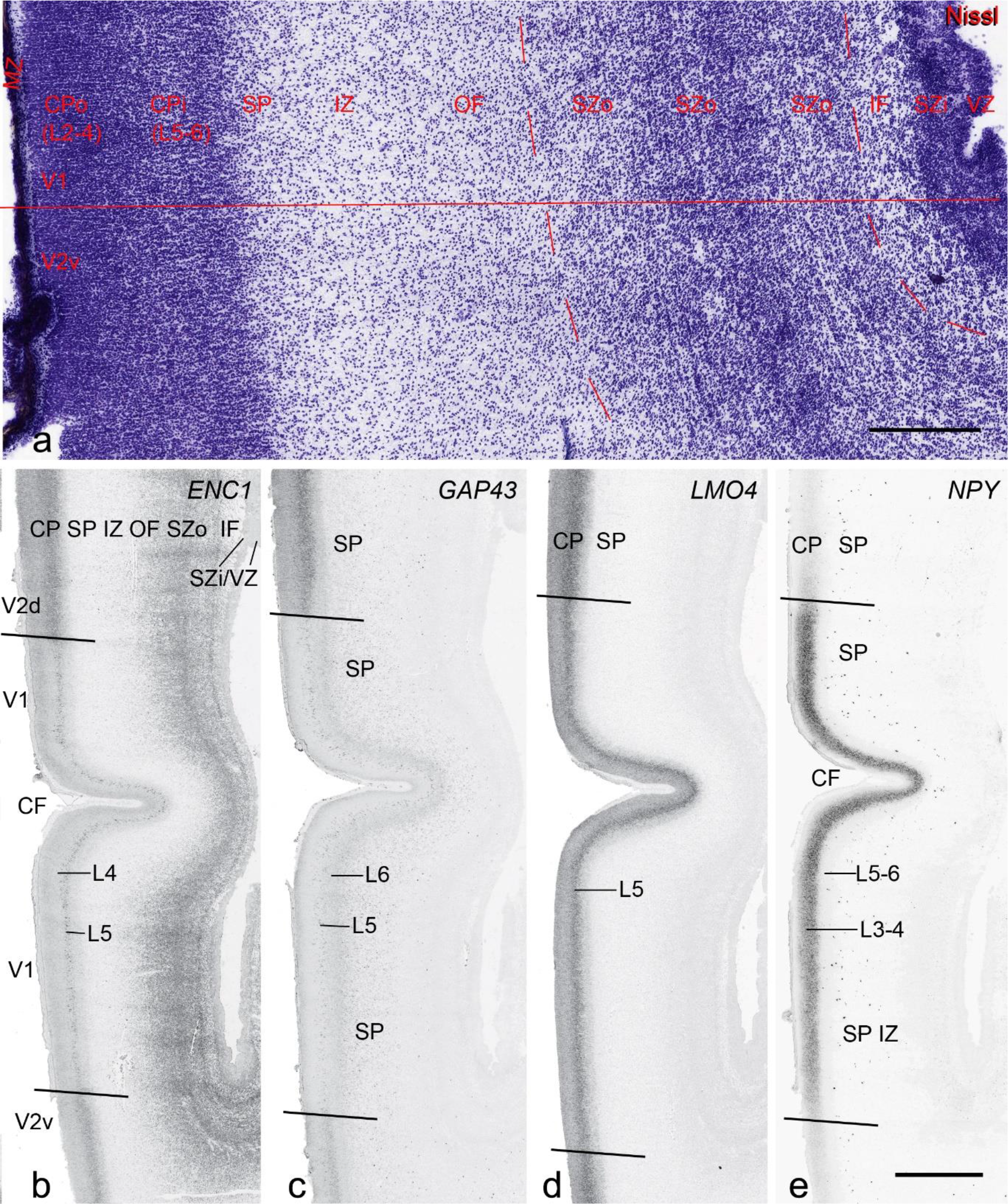
Cytoarchitecture and gene expression in the medial occipital cortex at PCW 21. (a). A Nissl-stained section showing the lamination of the cortex. CPo (future layers 2 to 4) and CPi (future layers 5 and 6) can be appreciated but differentiation between V1 and V2 (here V2v) is not yet clear in Nissl preparations. Inner fiber zone (IF) can be identified as a cell-less zone between SZi and SZo. (b-e). Expression patterns of *ENC1* (b), *GAP43* (c), *LMO4* (d) and *NPY*(e). Note the obvious difference of the gene expression patterns between V1 and the dorsal and ventral V2 (V2d and V2v, respectively). Scale bars: 400 µm in (a); 1600 µm in (b) for panels (b)-(e).

### 3.3 Delineation of prenatal neocortical areas

#### PCW 15

On Nissl-stained sections, obvious differences among neocortical regions were not observed at PCW 15. However, an anterior-posterior (A-P) gradient of gene expression in neocortex was observed. For instance, *LMO4* displays strong expression in the CP of frontal and temporal cortices with gradually weaker expression in parietal and occipital cortices (**Fig. 6a, b**). In contrast, *NPY* expression in the CP is stronger in temporal (**Fig. 6c**) and occipital cortices than in parietal (**Fig. 6c**) and frontal (**Fig. 6d**) cortices. *NPY* expression in the SP also shows regional difference with relatively stronger expression in posterior and lateral neocortex and weaker expression in anterior and medial neocortex (**Fig. 6c, d**). In addition, *NTRK2* shows weak expression in the SP and CPi of frontal neocortex but gradually stronger expression in parietal, temporal and occipital cortices, and is strongest in occipital neocortex (see **Appendix 2**). Anterior-posterior differences in *FABP7*, *LMO4*, *GRIK2* and *SATB2* expression in different layers also occur between dorsomedial frontal and occipital cortices (compare **Fig. 3** to **Fig. 4**). However, primary sensory cortices (V1, primary auditory A1, and somatosensory S1) and primary motor cortex (M1) are not distinguishable from adjoining areas at PCW 15. The cingulate cortex can be identified based on its differential expression patterns of genes such as *ETV1*, *ENC1* and *LMO4* from adjoining regions (see **Appendix 2**). Therefore, frontal, parietal, temporal, occipital and cingulate cortices can be roughly identified at PCW 15.

**Fig. 6.**
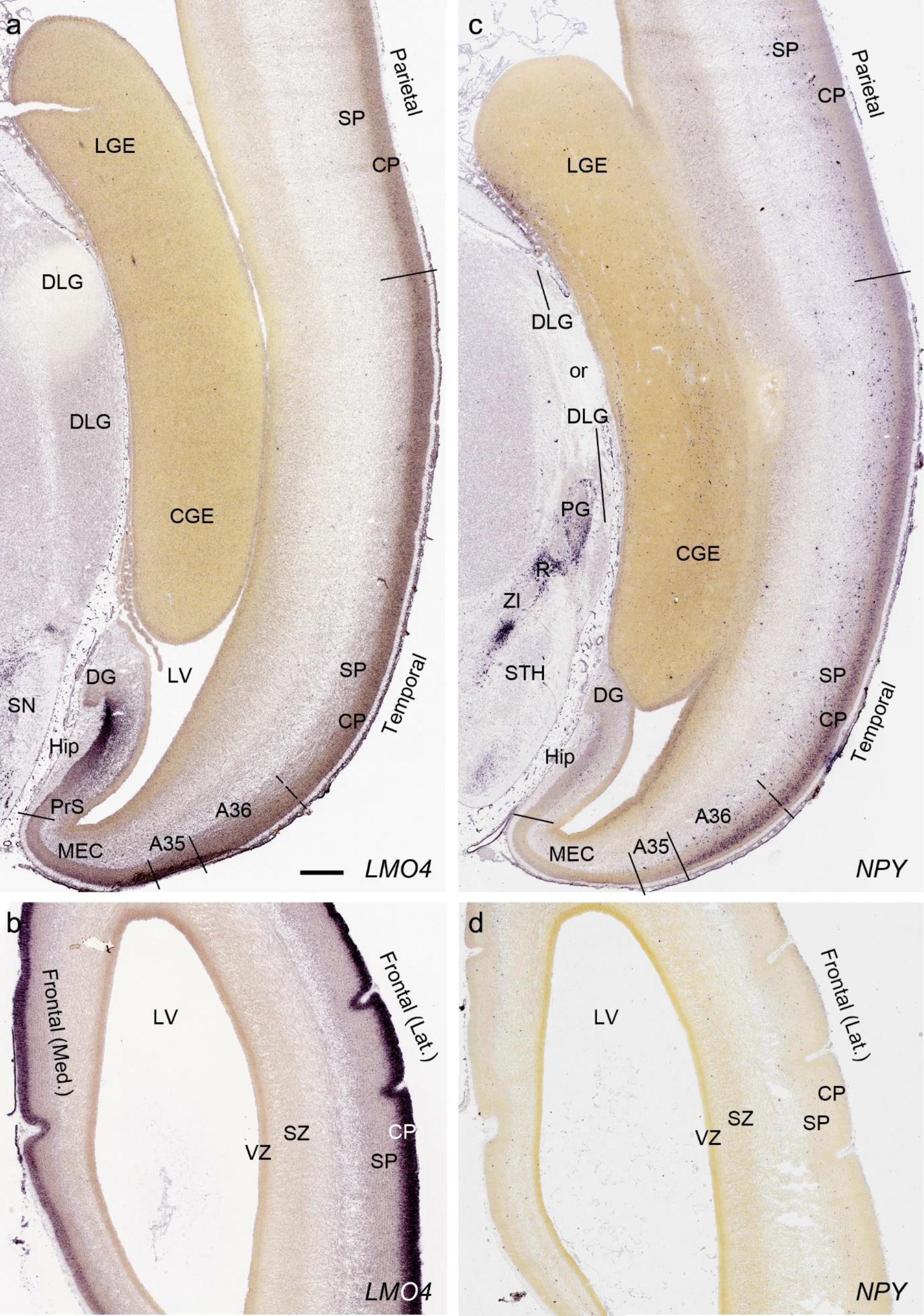
Differential gene expression across neocortex at PCW15. (a,b) expression of *LMO4* in parietal, temporal (a) and frontal (b) cortices. Note the strong expression in the hippocampus (Hip). (c,d) differential expression of *NPY* in parietal, temporal (c) and frontal (d) cortices. *NPY* is also expressed in the pregeniculate (PG) and reticular thalamic (R) nuclei. DLG, dorsal lateral geniculate nucleus; SN, substantia nigra; ZI, zona incerta; STH, subthalamic nucleus. Scale bar: 790 µm in (a) for all panels.

#### PCW 21

In addition to the identified major neocortical regions described above, V1 can be distinguished from the secondary visual cortex (V2) at PCW 21 on *ENC1-, GAP43-, LMO4-*and *NPY*-ISH sections (**Fig. 5b-e**). Generally, the former three genes are much less expressed in V1 than in V2 (**Fig. 5b-d**), while *NPY* is strongly expressed in the deep CP of V1 compared to V2 (**Fig. 5e**). However, A1 and S1 cannot be well distinguished from adjoining cortices. Subtle difference between M1 and S1 appears at PCW 21, for example, on *ENC1-, PLXNA2-, NRGN-*and *ETV1*-ISH sections. These gene markers clearly display layer 5 of the neocortex. As M1 has a well-developed and thicker layer 5 than S1, which shows a weaker layer 5, the border between M1 and S1 can be roughly established at PCW 21 (see **Appendix 4**). Similarly, the cingulate cortex can be identified at PCW 21 more easily than at PCW 15 based on the expression patterns of *ETV1*, *ENC1, LMO4, PLXNA2* and *NRGN* (see **Appendix 4**). For example, *ETV1* and *NRGN* are strongly expressed in both anterior and posterior cingulate cortex but only weakly in the adjoining neocortex (see **Appendix 4**). Finally, the dysgranular and granular insular cortex (Idg and Ig, respectively) can also be identified at PCW 21 based on Nissl stain, gene expression patterns, and its relationship with the claustrum, located deep to the insular cortex and displaying strong *GRIK2* and *LMO4* expression.

### 3.4 Delineation of the layers in prenatal allocortex and periallocortex

In contrast to the neocortex, which typically has six well-defined cortical layers, the allocortex, which includes the hippocampal formation (HF or archicortex) and olfactory cortices (mainly the piriform cortex, Pir), generally displays three major layers in mature cortex. As in the mature brain, the prenatal Pir is a three-layered, easily identified structure, and as such is not further described in this study. The HF in this study mainly contains the hippocampus [dentate gyrus (DG) and hippocampal subfields (CA1-4)] and the subicular cortex [prosubiculum (ProS), and subiculum (S)]. The cortical region located between allocortex and neocortex is usually termed periallocortex, which includes peripaleocortex and periarchicortex. The former mainly includes agranular insular cortex (Iag) and agranular liminal areas (areas FI and TI;) while the latter includes entorhinal cortex (EC), perirhinal cortex (area 35), presubiculum (PrS), parasubiculum (PaS) and retrosplenial cortex (RSC: areas 29 and 30) (see **Table 1** and Ding et al., 2016). The periallocortex has more than three layers (4 to 6 or 7 layers) and these layers are usually not equivalent to the neocortical layers. Note that other related terms were also used in literature. For example, subicular complex was used to include ProS, S, PrS and PaS (e.g. Ding, 2013). The medial temporal cortex (MTC) was used to contain PrS, PaS, EC and area 35 (similar to periarchicortex without RSC). The following description mainly focuses on the HF and MTC.

#### PCW15

On Nissl preparations, the typical laminar organization of the HF is obvious at this stage (**Fig. 7a**). Many genes show clear expression patterns in distinct layers or sublayers of the HF. The expression patterns of 16 genes are shown as examples (**Fig. 7b-p**). Specifically, the CP of the HF (the, hippocampal plate) expresses *FADS2*, *FOXP1*, *ETV1* and *SYNGAP1* in its full thickness, while the inner CP (CPi) expresses additional genes such as *FEZF2*, *NRTK2*, *NRGN* and *SHANK3*. *FABP7*, *FADS2* and *NTRK2* are strongly expressed in the VZi while strong expression of *VIM* is seen throughout the VZ. Interestingly, *TBR2* expression appears to concentrate at SZi/VZo border or deep SZ (SZd). The SP expresses *GRIK2*, while *GAP43*, *FOXG1* and *ENC1* are strongly expressed in the SZi, CP and SP, but weakly in IZ. In the MZ strong expression was found for *GAP43* (**Fig. 7c**), *ERBB4*, *DCX*, *RELN* and *CALB2* (see **Appendix 2**). Interestingly, SZo is not identified in the HF and MTC. As shown in **Figure 8**, only the thinner VZ and SZi, but not the thicker SZo, extend from temporal neocortex into the MTC and HF. The SZi is recognizable by a lower expression of *TBR2* (**Fig. 8a**) and *VIM* (**Fig. 8b**) compared to the VZ and SZo. The existence of SZi in the MTC and HF are also revealed by the strong expression of *TBR2* between the VZ and SZi, a zone sometimes termed as deep SZ (**Fig. 8a**).

**Fig. 7.**
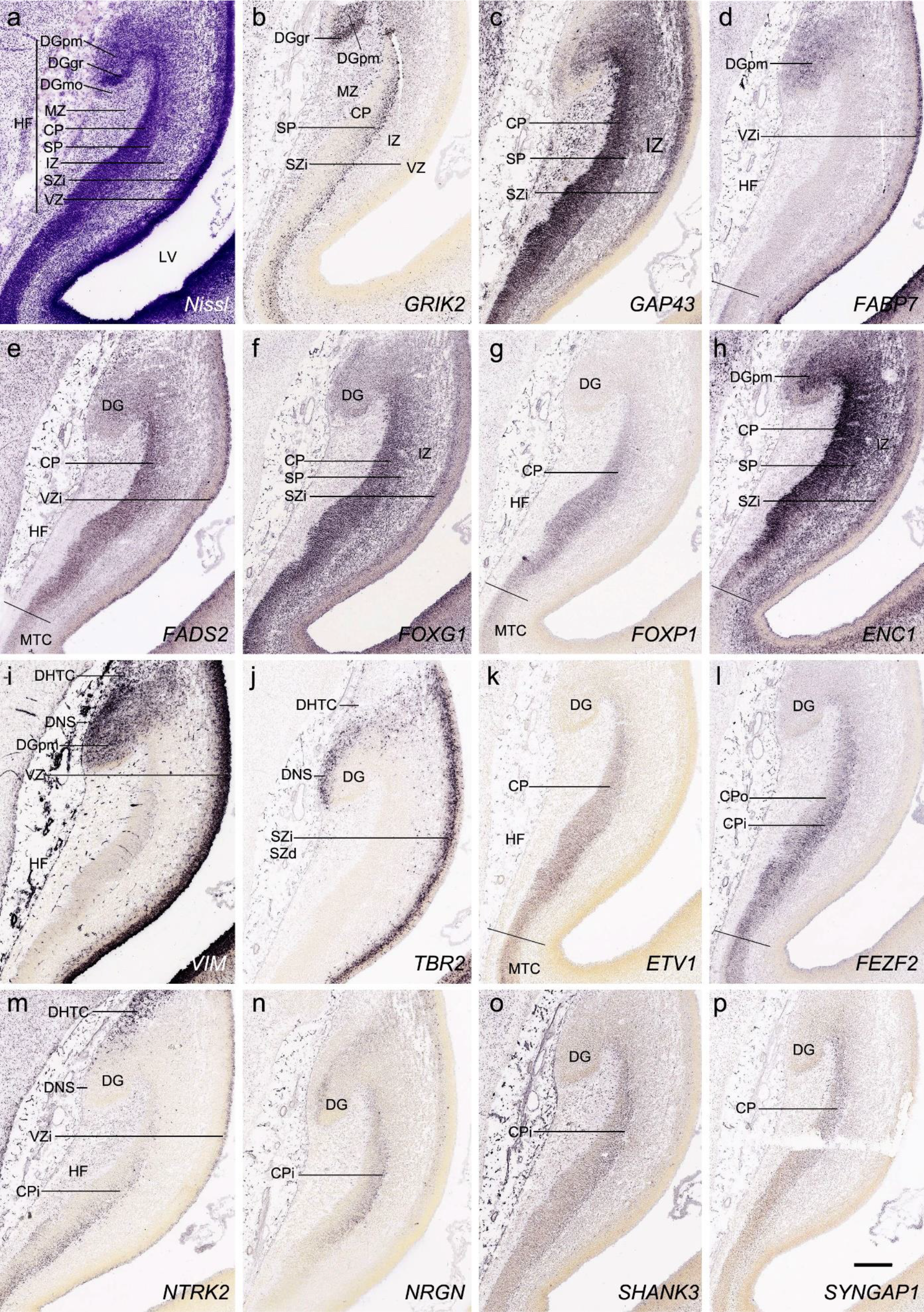
Gene expression in hippocampal formation (HF) at PCW15. (a). A Nissl-stained section showing the laminar organization of the HF. (b-p). Expression patterns of 15 genes as indicated in (b-p). Note the layer-specific gene expression in HF and the expression of *VIM* (i) and *NTRK2* (m) in the migrating dentate-hippocampal transient cells (DHTC). VZi, inner VZ; DG, dentate gyrus; DGmo, DGgc and DGpm, molecular, granular and polymorphic layers (zones) of the DG. Scale bar: 400 µm in (p) for all panels.

**Fig. 8.**
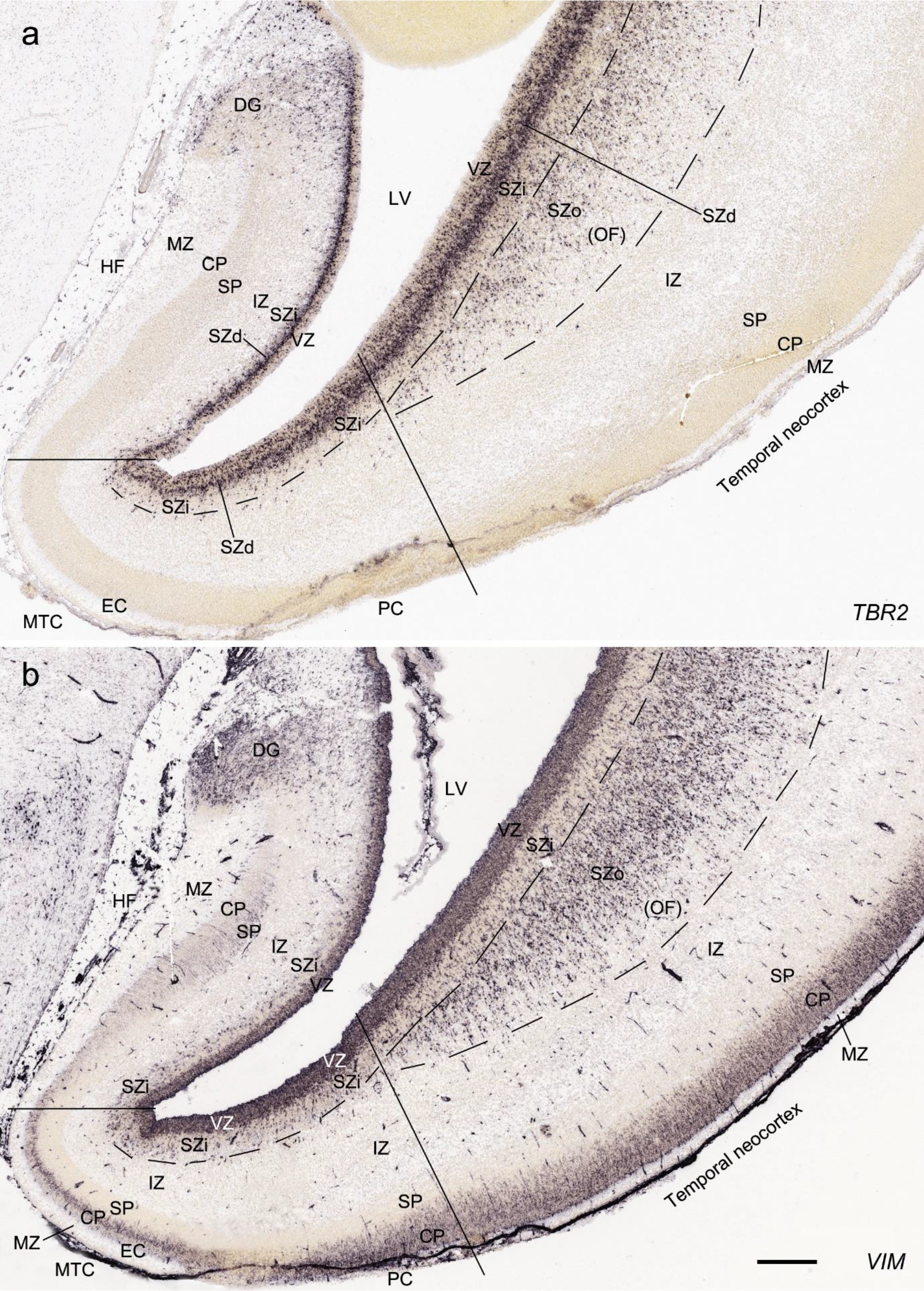
Comparison of the layers in allocortex, periallocortex and neocortex at PCW 15. *TBR2* (a) and *VIM* (b) are expressed in the VZ and SZi of these three types of cortex as well as in the SZo of the neocortex. Note that the SZo in the temporal neocortex does not extend into periallocortex [mainly entorhinal (EC) and perirhinal (PC) cortices] and hippocampal formation (HF). *VIM* is also strongly expressed in the CP of the neocortex and PC (i.e. area 35) but weakly in the CP of the EC (b). Scale bar: 400 µm in (b) for (a, b).

#### PCW 21

All the layers of the HF and MTC seen at PCW 15 can be identified on Nissl-stained and ISH sections at PCW 21. These layers include MZ, CP, SP, IZ, SZi and VZ (**Fig. 9a**). Layer-specific gene expression is also observed. *SOX2, FOXG1, ENC1, GAP43, NTRK2, SHANK3, SYNGAP1, LBX1, LHX2, NRGN, LMO4, DCX* (see **Appendix 4**) and *NES* (**Fig. 9a**) are strongly expressed in the CP while *FOXP1* and *CNTNAP2* expression (**Fig. 9c**) are strongly expressed in the inner CP (CPi). *RELN* is expressed in MZ (**Fig. 9b**), and *NPY, SST, PLXNA2* (see **Appendix 4**) and *GRIK2* (**Fig. 9d**) in SP. Finally, *PAX6* is lightly expressed in SZi (**Fig. 9e**) and *SOX2, VIM* (not shown) and *GFAP* strongly in VZ (**Fig. 9f**). The thick SZo does not extend from the temporal neocortex into the MTC and HF, as demonstrated using SZo markers such as *FABP7* (not shown), *NES, PAX6, TBR2* and *VIM* (**Figs. 9a, e; 10a, d**). In the MTC, *ETV1* and *NRXN1* are also mostly expressed in layers 5-6 of the EC (**Fig. 11a-c**). In contrast, *GRIK2* and *LMO4* are mainly expressed in layers 2-3 of the EC with relatively lower expression in L5-6 (**Fig. 11d, e**).

**Fig. 9.**
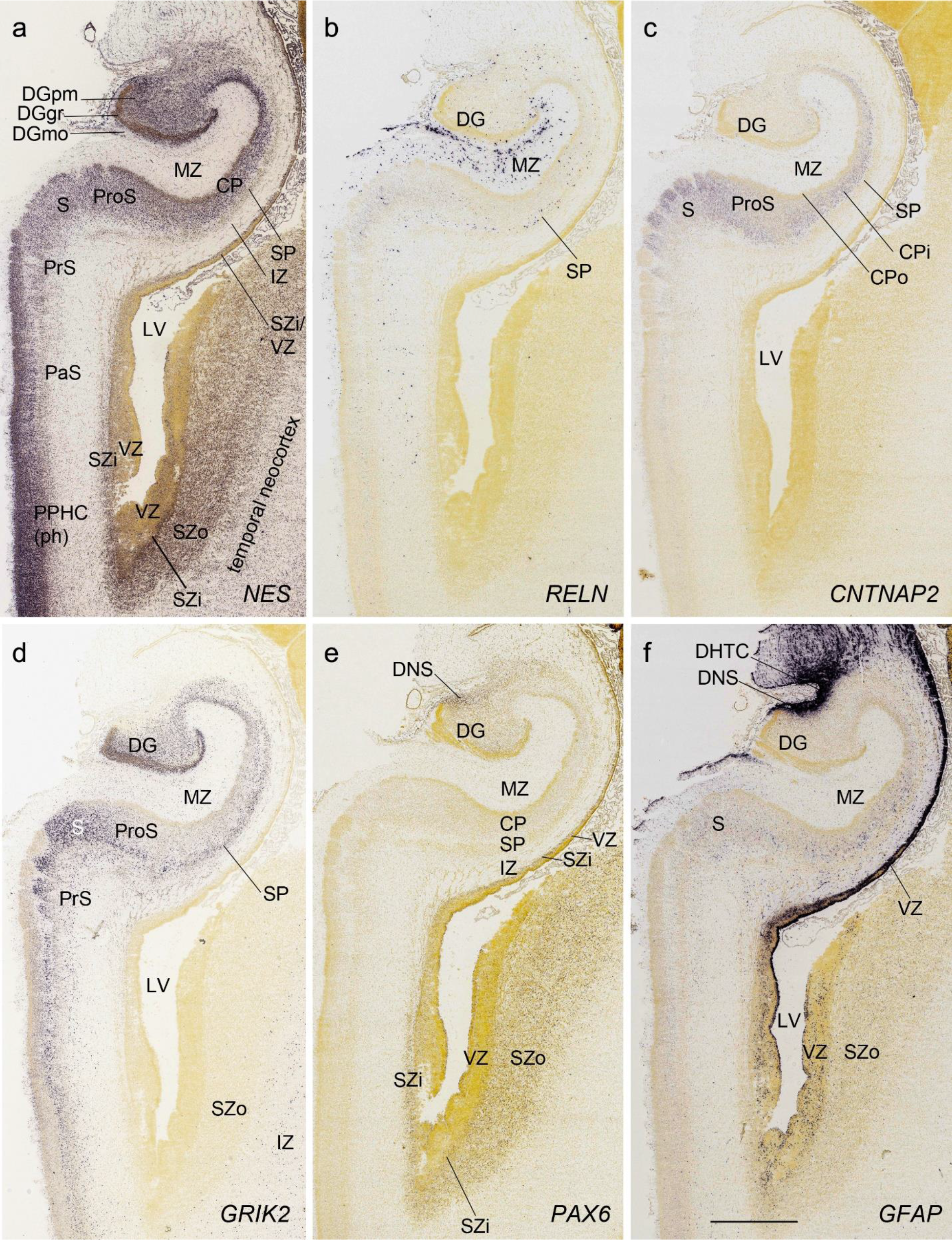
Layer-selective gene expression in the hippocampus at PCW 21. (a). *NES* expression in the CP but not the SP of the hippocampus. (b). *RELN* expression in the MZ and SP of the hippocampus and DGmo. (c, d). *CNTNAP2* (c) and *GRIK2* (d) expression in the inner CP (CPi) of the hippocampus and superficial layers of the subiculum (S). (e). *PAX6* expression in the SZi of the hippocampus and SZo of the temporal neocortex. (f). *GFAP* expression in the VZ of the hippocampus and DNS of the DG. Scale bar: 1590 µm in (f) for all panels.

**Fig. 11.**
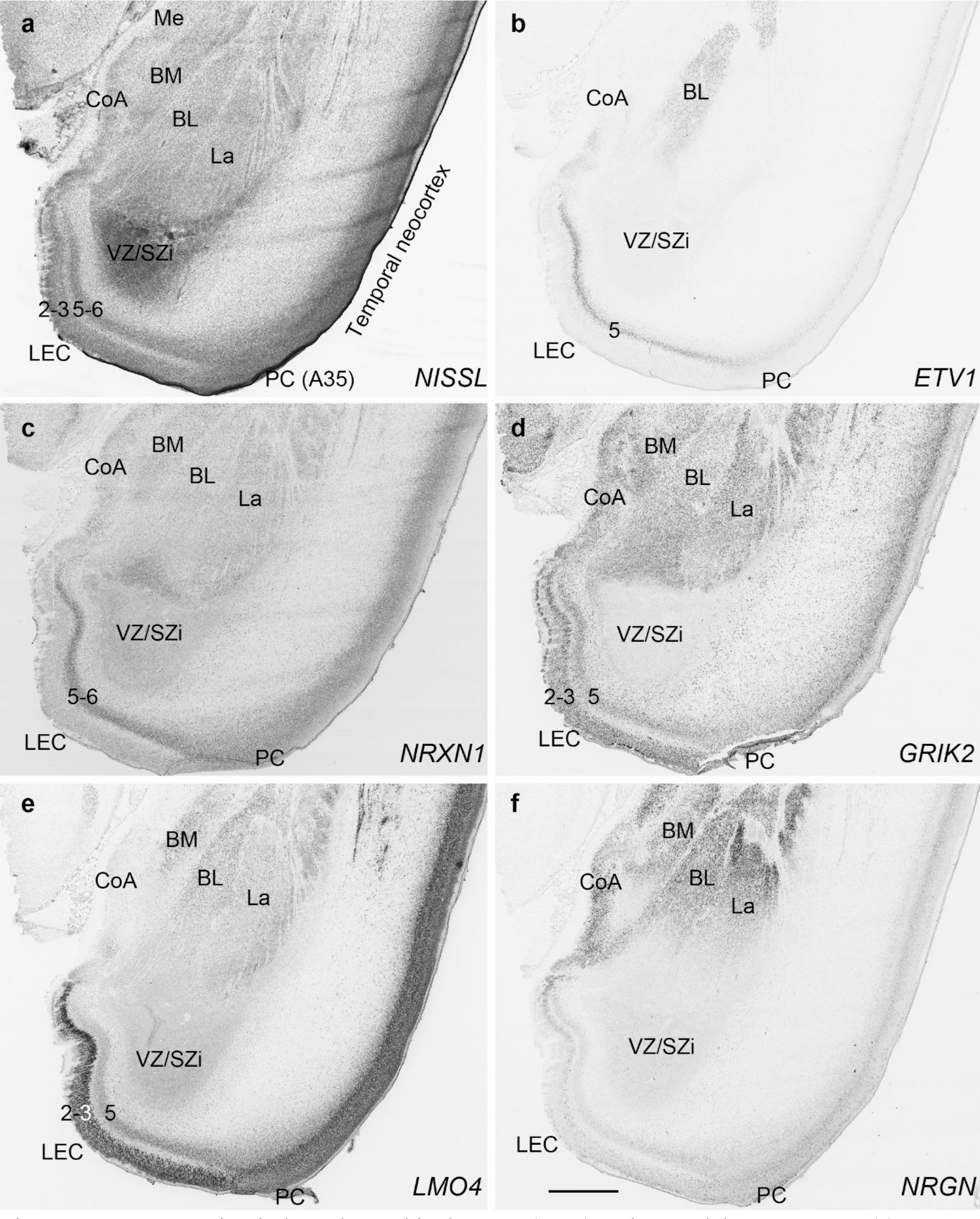
Gene expression in lateral entorhinal cortex (LEC) and amygdala at PCW 21. (a) A Nissl-stained section showing the cytoarchitecture in the LEC and amygdaloid nuclei. (b-f) Expression patterns of *ETV1* (b), *NRXN1* (c), *GRIK2* (d), *LMO4* (e) and *NRGN* (f) in the LEC and amygdaloid nuclei. Scale bar: 1590 µm in (f) for all panels.

In summary, a striking feature of the HF and MTC appears to be its lack of SZo, which is one of the thickest neocortical layers at PCW15 and 21. The thickness of SZo in temporal neocortex is dramatically reduced towards the border with the perirhinal cortex (**Fig. 8**) and the SZo is not observed in the MTC and HF (**Figs. 8; 9; 10**). At PCW 21, the VZ and SZi extend from the temporal neocortex into the HF with gradually narrowing of their thickness towards the DG (e.g., **Figs. 9a; 10a, d**).

### 3.5 Delineation of the subregions in prenatal allocortex and periallocortex

#### PCW 15

To investigate whether regional difference can be distinguished in the prenatal allocortex and periallocortex, we analyzed and compared gene expression patterns in different regions of the HF and MTC, and between the MTC and neocortex. At PCW 15, the granular layer of the DG (DGgr) is recognizable from the remaining HF (**Fig. 7a**, b). The polymorphic layer of the DG (DGpm) can also be roughly identified in *GRIK2, GAP43, FABP7,* and *ENC1*-ISH sections (**Fig. 7b, c, d, h**). However, subfields CA1-3 are not yet distinguishable within the HF although the boundary between the HF and MTC is appreciable (**Fig. 7d, e, g, h**, k, l). Interestingly, strong *VIM* and *TBR2* expression is observed in the so-called dentate neuroepithelial stem cell zone (DNS; see Nelson et al., 2020) while *VIM* and *NTRK2* are strongly expressed in another transient zone called dentate-hippocampal transient cell zone (DHTC), which is located immediately dorsal to the DNS (**Fig. 7i**, m). The boundaries between the MTC and neocortex are identifiable based on expression patterns of some genes in addition to the lack of SZo in the MTC. For instance, *SATB2* displays strong expression in the entire CP and IZ and moderate expression in the SP of the insular and temporal neocortex (**Fig. 12a**, b). In contrast, strong *SATB2* expression is only seen in the IZ and CPi (layers 5-6) of the lateral EC (LEC; **Fig. 12a**) and in layer 5 of the medial EC (MEC; **Fig. 12b** and the inset in b). In addition, *ETV1* is strongly expressed in the MTC but lightly in the neocortex at PCW 15 (see **Appendix 2**).

**Fig. 12.**
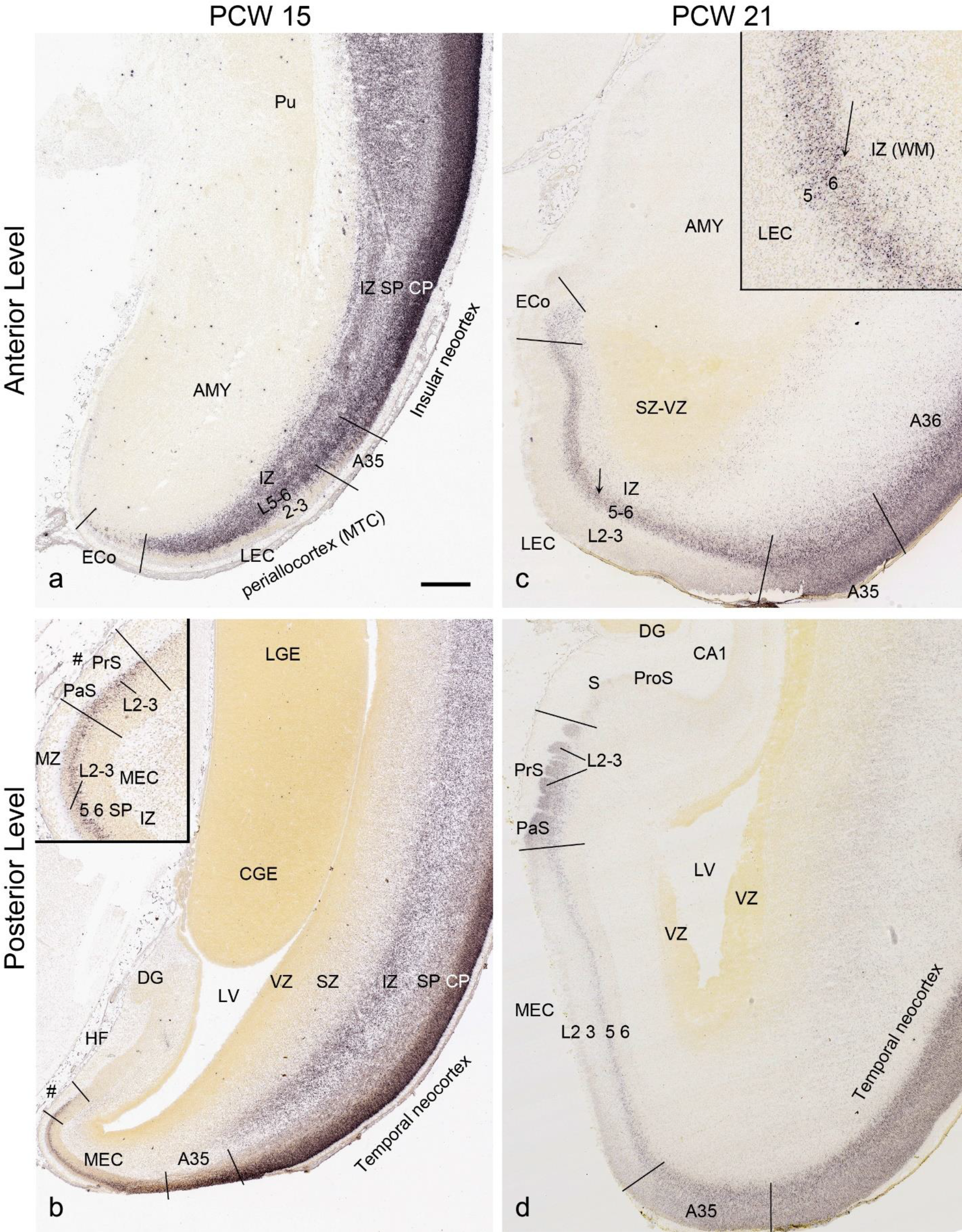
Identification of lateral and medial entorhinal cortex at PCW 15 and 21. (a, b) Lateral (a) and medial (b) entorhinal cortex (LEC and MEC, respectively) identified on *SATB2*-stained section at PCW15. The inset in (b) is a higher power view of the MEC (#s indicate the same location). (c, d) LEC (c) and MEC (d) identified on *SATB2*-stained section at PCW21. The inset in (c) is a higher power view of the LEC (the arrows indicate the same location). *SATB2* expression is seen in both layers 5 and 6 of the LEC including the olfactory part (ECo) (c) but only in layer 5 of the MEC (d). At both PCW 15 and 21, strong *SATB2* expression is seen in layer 5 of the MEC and layers 2-3 of the presubiculum (PrS) and parasubiculum (PaS). Pu, putamen; AMY, amygdala; LGE and CGE, lateral and caudal ganglion eminence. Scale bar: 790 µm in (a) for all panels.

#### PCW 21

Regional differences within the HF are clear at PCW 21. For instance, the subiculum strongly expresses *CNTNAP2, GRIK2* (**Fig. 9c**, d), *ETV1 and FEZF2* (**Fig. 10b**, c) while adjoining HF regions (e.g., ProS, CA1-3) display very low expression of these genes. Alternatively, some genes such as *TBR2* and *VIM* show strong expression in the DGpm and DNS but not or little expression in other HF regions (**Fig. 10a**, d). The DNS also strongly expresses *PAX 6, GFAP* (**Fig. 9e**, f), *SOX2,* and *NTRK2* while the DHTC express strong *GFAP*, weaker *VIM, SOX2* and *NTRK2*, and no or low *TBR2* and *PAX6* (see **Appendix 4**). The regional difference between MTC and neocortex at PCW 21 is even more obvious than that at PCW 15 (**Fig. 12c**, d). LEC and MEC have strong *SATB2* expression in layers 5-6 and layer 5, respectively (**Fig. 12c**, d). In the PrS and PaS, strong *SATB2* expression is seen in the superficial rather than deep layers (**Fig. 12d**). In contrast, no *SATB2* expression is observed the subiculum and hippocampus (**Fig. 12d**).

**Fig. 10.**
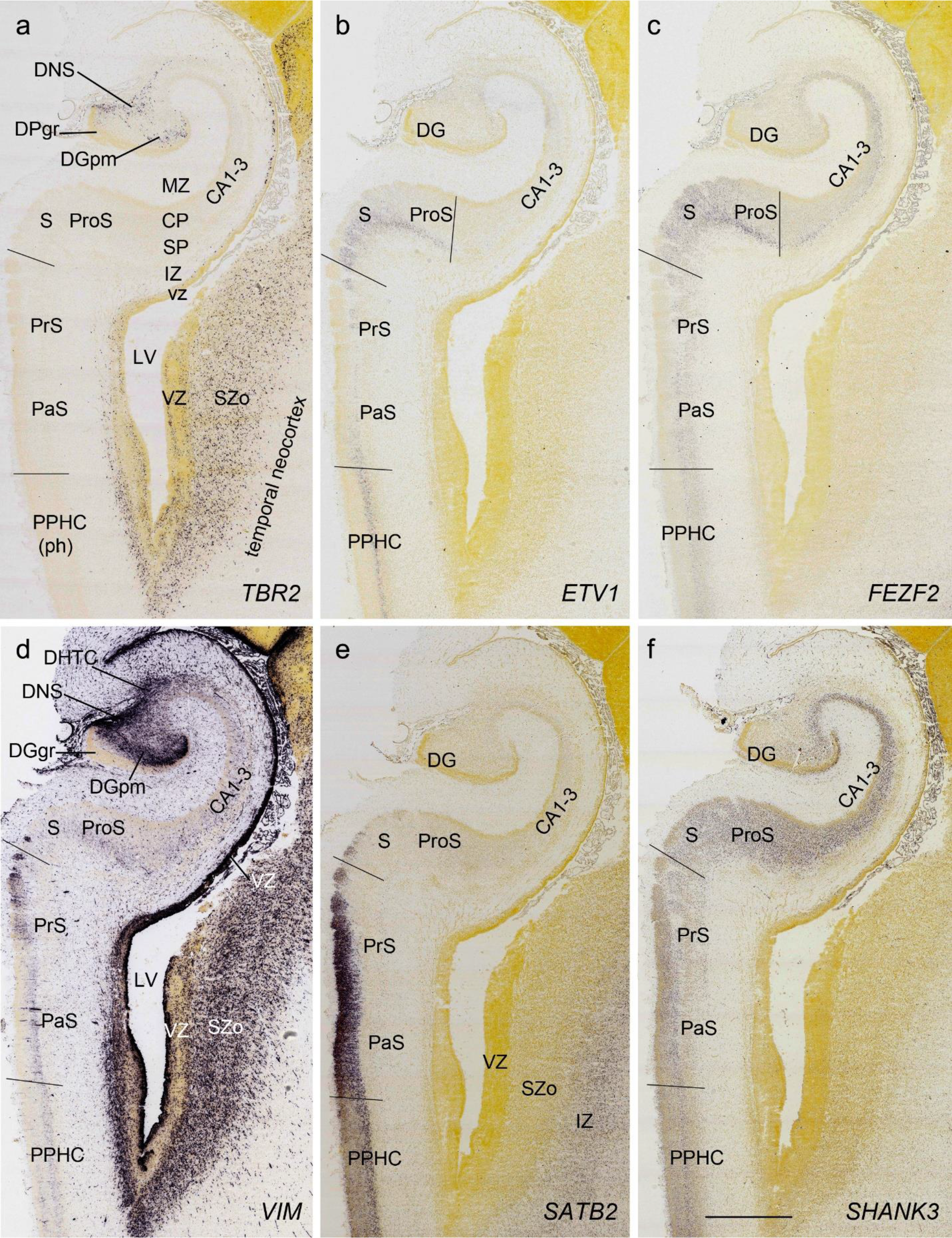
Region-selective gene expression in the HF at PCW 21. (a). *TBR2* expression in the DNS and DGpm. Strong *TBR2* expression is also seen in the SZo of the temporal neocortex. (b, c). *ETV1* (b) and *FEZF2* (c) expression in the subiculum (S) and deep prosubiculum (ProS). *FEZF2* is also expressed in the deep CP of the hippocampus (CA1-3). (d). Strong *VIM* (c) expression in DNS, DHTC, DGpm and VZ of the hippocampus as well as in the VZ and SZo of the temporal neocortex. (e). Strong *SATB2* expression in the presubiculum (PrS) and parasubiculum (PaS) as well as in the IZ of the temporal neocortex. (f). Strong *SHANK3* expression in the CP of the hippocampus. Scale bar: 1590 µm in (f) for all panels.

### 3.6 Delineation of cerebral nuclei

#### PCW 15

Strong gene expression was found in specific cerebral nuclei and these are helpful in their delineation. For example, *CALB2* expression is observed in the medial (Me), anterior cortical (CoA) and parts of the lateral (La) nuclei of the amygdala (**Fig. 13a**). Other such regional expression patterns are *NPY, SST, FOXG1* and *DLX1* in the bed nucleus of terminalis (BNST); *ETV1* expression in the external part of globus pallidus (GPe) and basolateral nucleus (BL) of the amygdala; *SST* expression in central nucleus (CEN) of the amygdala; *FOXP1* and *NPY* expression in caudate nucleus (Ca) and putamen (Pu); *GRIK2* and *LMO4* expression in claustrum; *NKX2.1* and *ZIC1* expression in GPe; and *RELN* and *LMO4* expression in GPi. In addition, *ZIC1* is strongly expressed in septal nucleus (SEP) and basal nucleus of Meynert (BNM). It should also be pointed out that strong transient expression of *CALB2* (**Fig. 13a**) and *PAX6* is observed in the space between the GPe and GPi. We refer to this zone as inter-pallidal transient cell zone (IPTC; **Fig. 13a**). *CALB2* is also expressed in the transient cells located between GPe and Pu (**Fig. 13a**). Expression of all above mentioned genes is shown in **Appendix 2**.

**Fig. 13.**
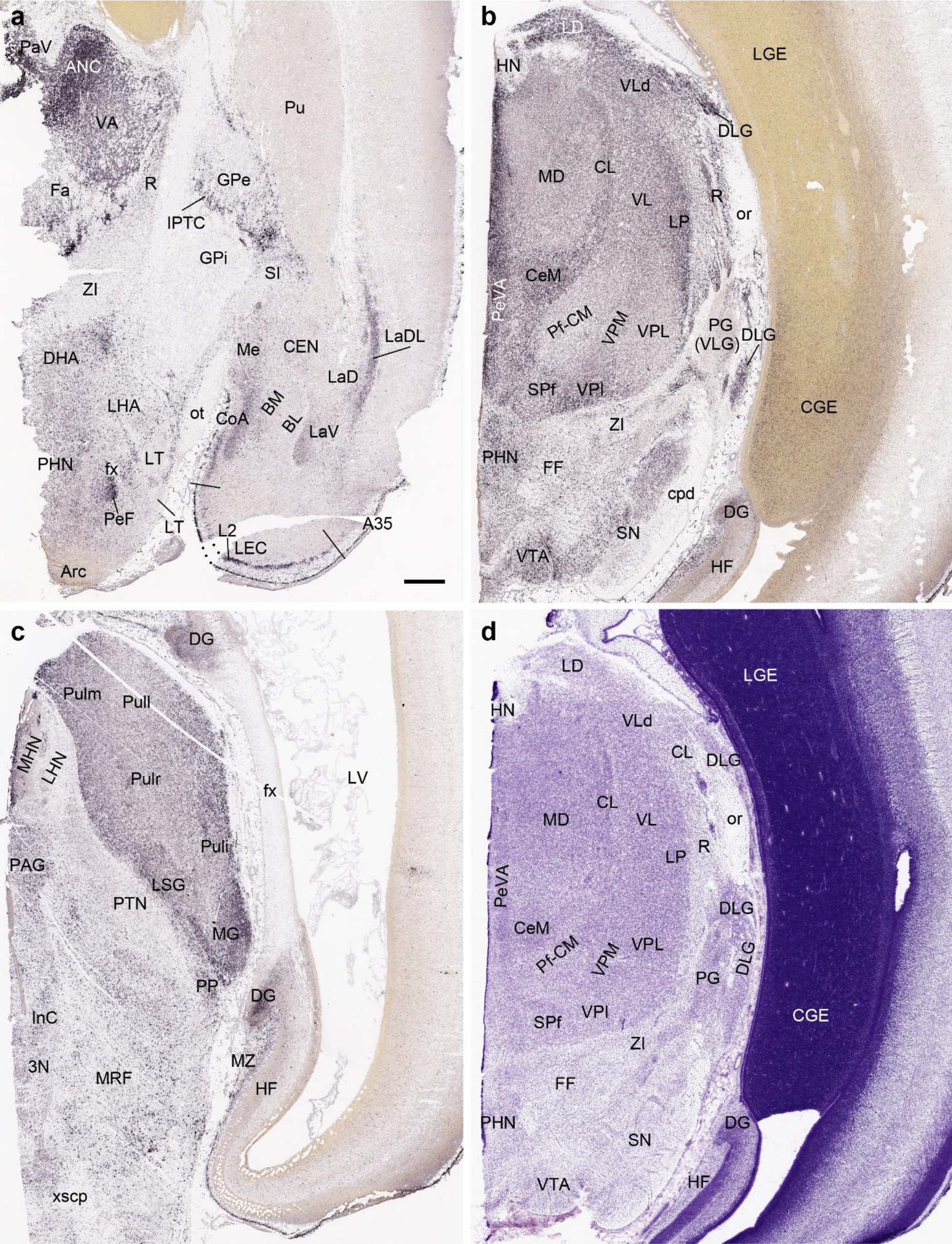
*CALB2* expression in the brain at PCW 15. (a) *CALB2* expression in the anterior thalamic region, basal ganglion, amygdala and lateral entorhinal cortex (LEC). Note the strong *CALB2* expression in anterior thalamic nuclear complex (ANC), medial hypothalamic region, anterior cortical nucleus of the amygdala (CoA), inter-pallidal transient cell zone (IPTC) and layer 2 of the LEC. (b) *CALB2* expression in middle thalamic region, midbrain and caudal ganglion eminence (CGE). Strong *CALB2* expression is seen in the lateral dorsal nucleus (LD), ventral lateral nucleus (VL), dorsal lateral geniculate nucleus (DLG), central medial nucleus (CeM) and subparafascicular nucleus (SPf) of the thalamus as well as in the ventral tegmental area (VTA) and CGE. (c). *CALB2* expression posterior thalamic region, midbrain and HF. Note the strong *CALB2* expression in the pulvinar (Pul), medial geniculate nucleus (MG), limitans/suprageniculate nucleus (LSG), medial habenular nucleus (MHN) and the dentate gyrus (DG). (d) A Nissl-stained section adjacent to (b) showing the cytoarchitecture of the thalamic regions. It is not easy to identify the different thalamic nuclei in Nissl-stained sections. In contrast, this is much easier in the *CALB2* ISH section (b). Scale bar: 800 µm in (a) for all panels.

#### PCW 21

In Nissl preparations, the major subdivisions of the amygdala can be identified (**Fig. 11a**). In ISH sections, *ETV1* is mostly expressed in the BL, while *NRXN1, GRIK2, LMO4* and *NRGN* are expressed in the BM, BL and La with various intensity in each subdivision (**Fig. 11c-f**). Strong *GRIK2* and *NRGN* expression is also observed in the anterior cortical nucleus of the amygdala (CoA; **Fig. 11d, f**). Gene expression patterns in other cerebral nuclei are similar to those at PCW 15 (see above).

### 3.7 Delineation of the thalamic and hypothalamic nuclei

Delineation of the thalamic nuclei at PCW 15 and 21 is mainly based on multiple region-specific gene markers such as *CALB2* (**Figs. 13, 14**), *CNTNAP2, PLXNA2, GRIK2, ETV1, NTRK2, ZIC1* and *ENC1* (see **Appendices 2** and **4**) as well as AChE staining (**Fig. 15**). *CALB2* is expressed in many thalamic nuclei including paraventricular (PaV), central medial (CeM), central lateral (CL), lateral dorsal (LD), ventral anterior (VA), ventral lateral (VL), reticular (R), subparafascicular (SPf), medial geniculate (MG), limitans-suprageniculate (LSG) and lateral posterior (LP) nuclei, as well as anterior nuclear complex (ANC; mainly AV and AM), pulvinar (Pul) and DLG (**Figs. 13a-c; 14a-g**). Strong expression of *CNTNAP2* occurs in mediodorsal (MD), centromedian (CM), dorsal lateral geniculate (DLG) and habenular (HN) nuclei of the thalamus, and *PLXNA2* expression in CM, parafascicular (Pf) and ventral posterior lateral (VPL) nuclei. In addition, *ZIC1* is a reliable marker for ANC, PaV, DLG, LD, Pul and medial HN (MHN) while *NRGN, SOX2, ZIC1* and *ETV1* are strongly expressed in MHN (see **Appendices 2** and **4**). As compared to **Figure 13b and 13d**, gene expression patterns are more helpful in the delineation of the thalamic nuclei than Nissl. For instance, strong and weak *CALB2* expression respectively in DLG and PG (VLG) allows an easy differentiation of the two structures (**Fig. 13b**) whereas this is difficult in Nissl-stained sections (**Fig. 13d**). Interestingly, we observe a region that is the likely equivalent of the mouse posterior intralaminar nucleus (PIL; Wang et al., 2020) in terms of its topographical relationship with adjoining regions and molecular signature. As in mouse, the PIL adjoins MG, SPf, posterior thalamic nucleus (Po), VPM, LSG, peripeduncular nucleus (PP) and pretectal nucleus (PTN) and has strong expression of *CALB2, GRIK2*, *LHX2 NRGN* and *NTRK2* (see **Appendices 2** and **4)**. Finally, it should be mentioned that AChE is a useful marker for the identification of the MD at PCW 15 (**Fig. 15a**) and 21 (**Fig. 15b, c**) and of the ventral posterior medial nucleus (VPM) and its parvocellular part (VPMpc) at PCW21 (**Fig. 15c**).

**Fig. 14.**
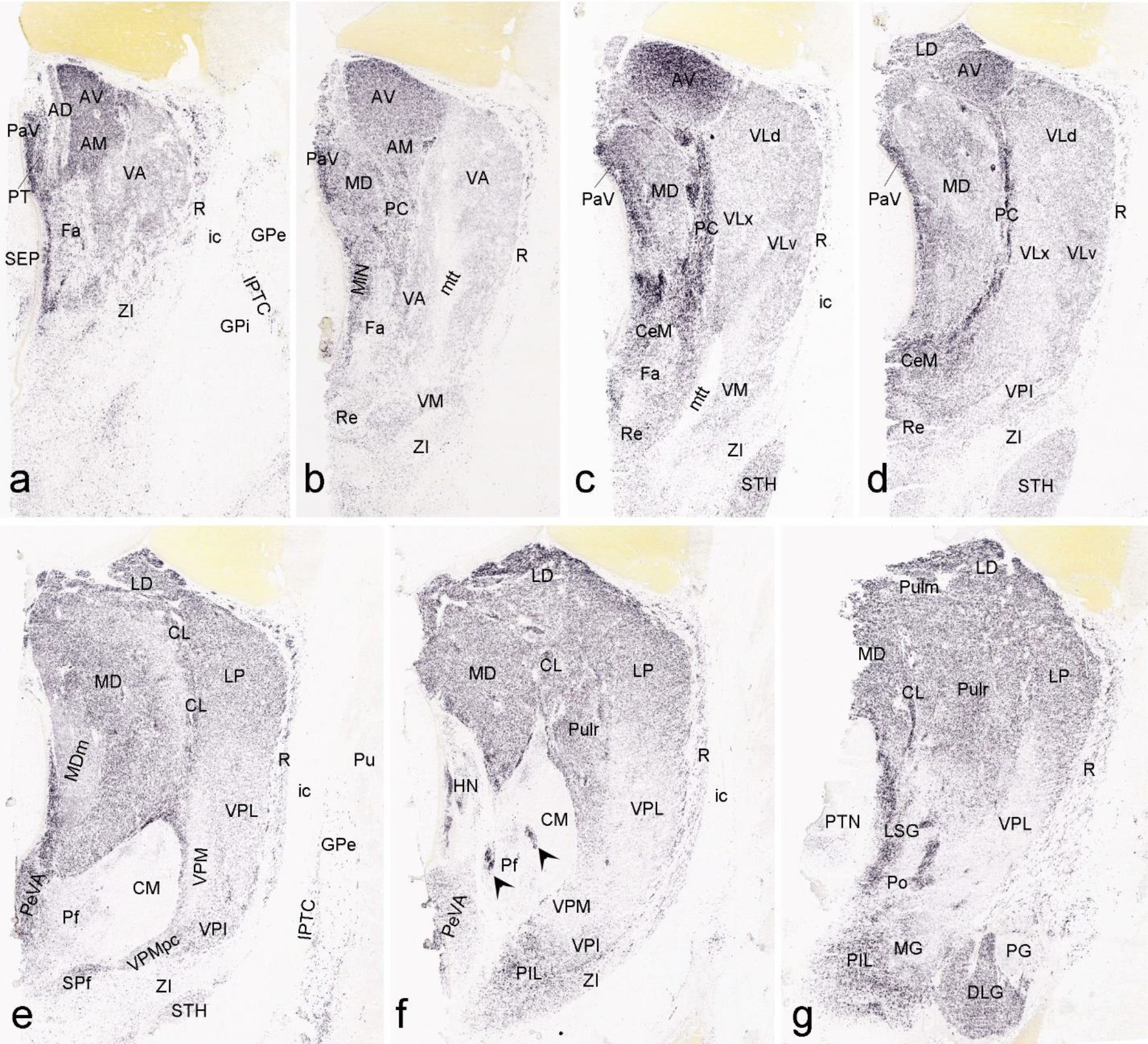
*CALB2* expression in the thalamus at PCW 21. (a-g) Sequential sections showing *CALB2* expression in the anterior ventral (AV) and anteromedial (AM) thalamic nuclei, paraventricular nucleus (PaV), midline nuclear complex (MiN), ventral anterior nucleus (VA), ventral lateral nucleus (VL), centromedian nucleus (CeM), reuniens nucleus (Re), and paracentral nucleus (PC) (a-d). Clear *CALB2* expression is also seen in the mediodorsal nucleus (MD), ventromedial nucleus (VM), lateral dorsal nucleus (LD), central lateral nucleus (CL), habenular nucleus (HN), periventricular area (PeVA), subparafascicular nucleus (SPf), lateral posterior nucleus (LP), pulvinar (Pul), dorsal lateral geniculate nucleus (DLG) and limitans/suprageniculate nuclei (LSG) (c-g). In contrast, faint *CALB2* expression is observed in ventral posterior lateral (VPL), ventral posterior medial (VPM), parafascicular nucleus (Pf), central medial nucleus (CM), pregeniculate nucleus (PG) and zona incerta (ZI) (e-f). Note the *CALB2* expression in the patches within Pf/CM (f). Note also the *CALB2* expression in the subthalamic nucleus (STH; c-e) and inter-pallidal transient cell zone (IPTC; a). Strong *CALB2* expression is also found in a region that appears to be equivalent to mouse posterior intralaminar nucleus (PIL). Scale bar: 800 µm in (a) for all panels.

**Fig. 15.**
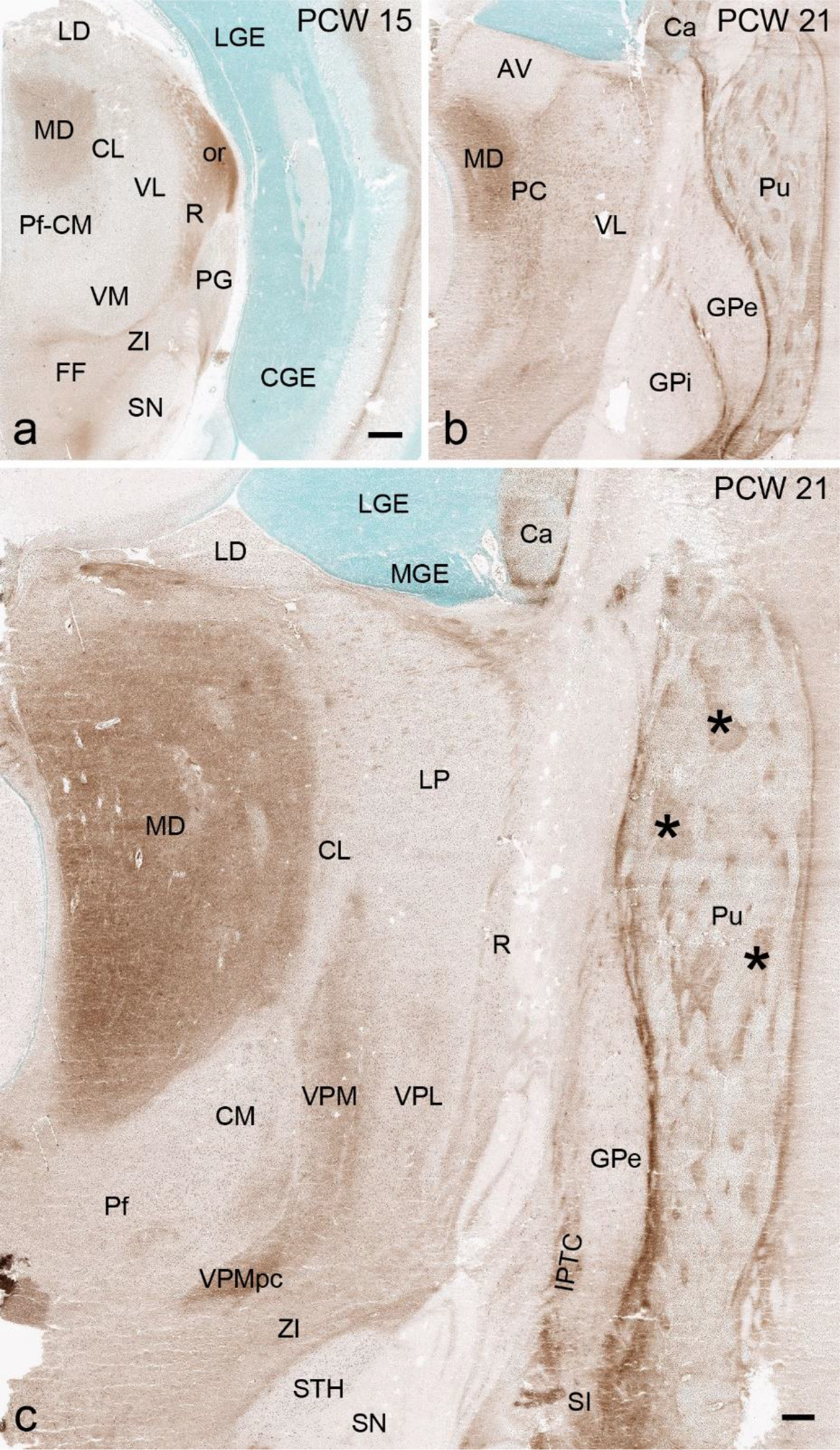
AChE staining patterns in the thalamus and basal ganglia. (a). AChE staining showing its enriched expression in the mediodorsal thalamic nucleus (MD) and some fiber regions such as optic radiation (or) at PCW 15. (b,c) AChE staining showing its enriched expression in the MD and two subdivisions of ventroposterior medial nucleus (VPM and VPMpc) as well as in the patches (*) of the putamen (Pu), substantia innominata (SI) and some fiber regions such as inter-pallidal transient cell zone (IPTC). Scale bars: 800 µm in (a) for (a-b); 400 µm in (c).

Genes with regional expression in hypothalamus at PCW 15 and 21 includes *CALB2*, which has strong expression in the ventromedial nucleus (VMH), perifascicular nucleus (PeF) and posterior hypothalamic nucleus (PHN) (**Fig. 13a, b**), and *SST*, which is expressed in the dorsomedial nucleus (DMH), paraventricular nucleus (PV) and lateral hypothalamic region (LH) (see **Appendices 2** and **4**). Other genes with region-specific expression include *GRIK2* in DMH; *DLX1* in anterior hypothalamic nucleus (AHN); *ZIC1* in suprachiasmatic nucleus (SCN) and PeF; *FOXG1* in medial preoptic nucleus (MPN); *LMO4* and *RELN* in the PV; *NKX2.1, CDH4* and *NTRK2* in VMH, and *NRXN1* in PHN and arcuate nucleus (Arc) (see **Appendices 2** and **4**).

### 3.8 Delineation of cerebellum and major brainstem structures

Cerebellar cortex at PCW 15 displays an immature shape with clear and thick external granular layer (EGL) and upper rhombic lip (URL) superficially, while deep cerebellar nuclei (CbDN) have formed in the center of the cerebellum (**Fig. 16a-j**). The EGL has strong expression of *PAX6* (**Fig. 16b, c**) and *ZIC1* while *ERBB4* and *GRIK2* are mainly expressed underneath the EGL (**Fig. 16h**). The URL contains strong *ZIC1* and *TBR2* expression (**Fig. 16j**). CbDN shows strong expression of *GAP43, CALB2, NRNX1, ERBB4* in the fastigial (Fas), interpositus (InP) and dentate (DT) nuclei (**Fig. 16e-h**), while *GRIK2* is mainly expressed in DT (**Fig. 16i**). These results are consistent with patterns observed in the mouse (e.g. **Fig. 15n-q**). It should be noted that *RELN* is not expressed in CbDN but present in surrounding cerebellar regions (**Fig. 17e, f**). In addition, some genes are mostly expressed in EGL of the vermis (e.g. *SST*) while others are strongly expressed in transient Purkinje cell clusters (TPC) (e.g., *CNTNAP2* and *NRXN1*; see **Appendix 2**).

**Fig. 16.**
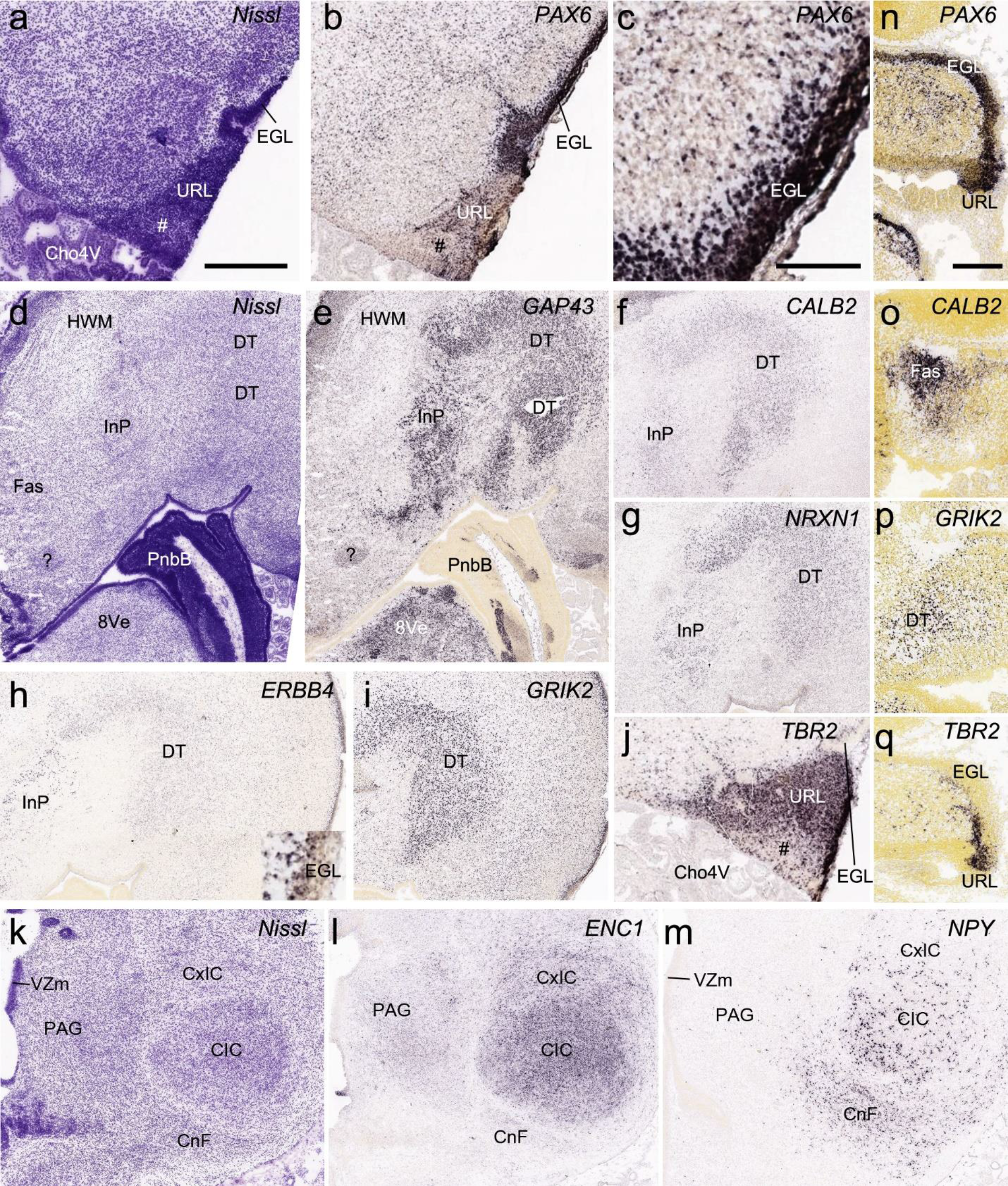
Cytoarchitecture and gene expression of the cerebellum and inferior colliculus. (a-l) from human brain at PCW 15; (n-q) from mouse brain at E15.5. (a) A Nissl-stain section showing the cytoarchitecture of the upper rhombic lip (URL) and external granular layer (EGL). This sectioning level is at about the level 40 of the atlas plates (see Appendix 1). (b, c) Low (b) and higher (c) power views of the *PAX6* expression in EGL. (d) A Nissl-stain section showing the cytoarchitecture of the cerebellar deep nuclei (CbDN), which include fastigial nucleus (Fas), interpositus nucleus (InP) and dentate nucleus (DT), and the pontobulbar body (PnbB), which is located at the junction of the cerebellum and pons. (e-i). Expression of *GAP43* (e), *CALB2* (f), *NRXN1* (g), *ERBB4* (h) and *GRIK2* (i) in the CbDN. Note that *ERBB4* and *GRIK2* are also strongly expressed in the region underneath the EGL (h, i). The inset in (h) is a higher power view of the EGL. (j) *TBR2* expression in the URL. The URL appears to have two parts with the less *TBR2 expression* part indicated by #. (K) A Nissl-stain section showing the cytoarchitecture of inferior colliculus (IC) and periaqueduct gray (PAG). (l, m) Expression of *ENC1* (l) and *NPY* (m) in the IC regions. (n-q) comparative expression of *PAX6* (n), *CALB2* (o), *GRIK2* (p) and *TBR2* (q) in the cerebellum at E15.5 (on sagittal sections). Scale bars: 400 µm in (a) for (a-m; except c); 100 µm in (c); 220 µm in (n) for (n-q).

**Fig. 17.**
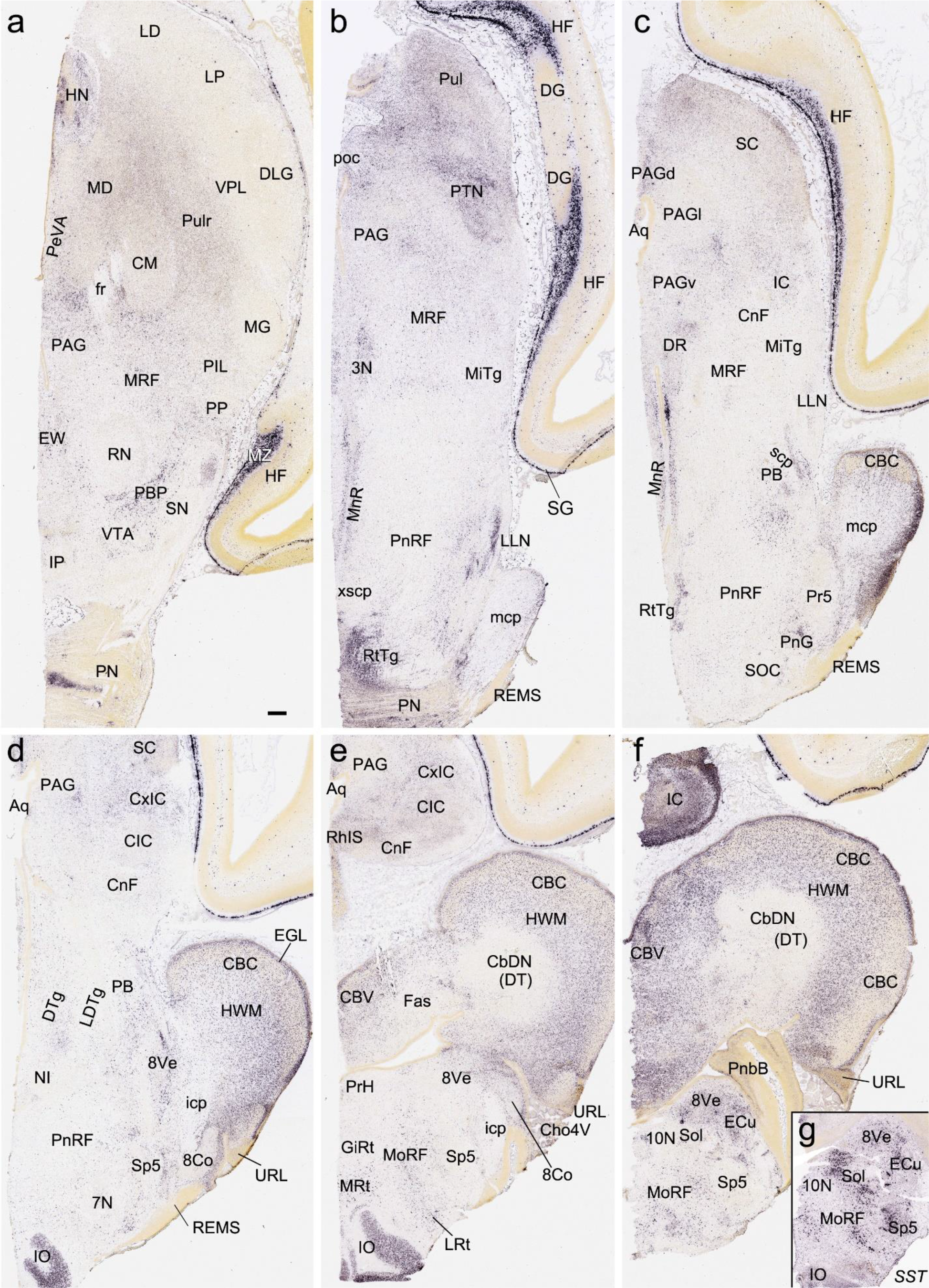
Gene expression in the brainstem and cerebellum at PCW 15. (a-f) *RELN* expression in brainstem nuclei and cerebellar cortex (CBC). Note the negative expression in cerebellar deep nuclei (CbDN) and strong expression in the hindbrain white matter (HWM). Note also the strong *RELN* expression in marginal zone (MZ in a) of the hippocampus and subpial granular zone (SG in b) of the cortex. (g) *SST* expression in the medullar nuclei. Scale bars: 400 µm in (a) for all panels. **Abbreviations:** 3N, oculomotor nucleus; 7N, facial nucleus; 8Co, cochlear nucleus; 8Ve, vestibular nucleus; 10N, vagal nucleus; Aq, aqueduct; CBV, cerebellar vermis; Cho4V, choroid plexus of 4^th^ ventricle; CM, centromedial nucleus; CnF, cuneiform nucleus; CxIC, cortex of inferior colliculus; DR, dorsal raphe nucleus; DTg, dorsal tegmental nucleus; ECu, external cuneate nucleus; EW, Edinger-Westphal nucleus; GiRt, gigantocellular reticular nucleus; HN, habenular nucleus; icp, inferior cerebellar peduncle; IO, inferior olive; IP, interpeduncular nucleus; LDTg, dorsolateral tegmental nucleus; LLN, nucleus of lateral lemniscus; LP, lateroposterior nucleus; LRt, lateral reticular nucleus; mcp, middle cerebellar peduncle; MiTg, microcellular tegmental nucleus; MnR, median raphe nucleus; MoRF, medullar reticular formation; MRF, midbrain reticular formation; MRt, medial reticular nucleus; NI, nucleus incertus; PB, parabrachial nucleus; PBP, parabrachial pigmented nucleus; PeVA, periventricular area; PIL, posterior intralaminar nucleus; PN, pontine nucleus; PnG, pontine gamma nucleus; PnRF, pontine reticular formation; poc, posterior commissure; PP, peripeduncular nucleus; Pr5, principal sensory nucleus of the trigeminal nerve; PrH, prepositus hypoglossal nucleus; PTN, pretectal nucleus; Pulr, rostral pulvinar; REMS, rostral extramural migration system; RhIS, rhombencephalic isthmus; RN, red nucleus; RtTg, reticular tegmental nucleus; scp, superior cerebellar peduncle; SN, substantia nigra; SOC, superior olivary complex; Sol, solitary nucleus; Sp5, spinal trigeminal nucleus; VPL, ventroposterior lateral nucleus; xscp, decussation of superior cerebellar peduncle.

In the midbrain, a rough lamination in the superior colliculus (SC) has formed at PCW15 with *FOXP1* and *VIM* expressed in the inferior gray layer (InG). *VIM* is also expressed in the periaqueductal gray region (PAG) (see **Appendix 2**). In the inferior colliculus (IC), central IC nucleus (CIC; **Fig. 16k**) has strong expression of *ENC1* (**Fig. 16l**), *NRGN* and *NPY*, the latter being also strongly expressed in the cuneiform nucleus (CnF; **Fig. 16m**). The cortical (or external) IC (CxIC) displays expression of *RELN* (**Fig. 17c-e**). The red nucleus (RN) shows strong expression of *CNTNAP2, GRIK2* and *NTRK2,* while the substantia nigra (SN) strongly expresses *ENC1* and *FABP7*. In addition, the parabrachial pigmented nucleus (PBP) located between RN and SN has strong *RELN* expression (**Fig. 17a**). *RELN* is also strongly expressed in the median raphe nucleus (MnR) and the oculomotor nucleus (3N) (**Fig. 17b, c**). In the lower brainstem (pons and medulla), *ZIC1, RELN* and *SST* are strongly expressed in the vestibular nuclei (8Ve) and external cuneate nucleus (ECu) (**Fig. 17f, g**) while *ENC1, ETV1, FOXP1* and *PLXNA2* (see **Appendix 2**) are expressed strongly in the inferior olive (IO). *RELN* is strongly expressed in the reticulotegmental nucleus (RtTg), lateral lemniscus nucleus (LLN), IO, and medullary reticular formation (MoRF) (**Fig. 17b, d, e, f**). *VIM* expression is found in 3N, Edinger-Westphal nucleus (EW) and abducens nucleus (6N) (see **Appendix 2**). Finally, the pretectal nuclear complex (PTN) contains strong expression of *RELN* (Fig. 16b), *FABP7* and *NPY* (see **Appendix 2**).

### 3.9 Delineation of the ganglionic eminence subdivisions

The ganglionic eminence (GE) mainly consists of three major parts, MGE, LGE and CGE (Hansen et al., 2013; Ma et al., 2013). In this study, the boundary between GE and adjoining cortical regions can be clearly identified with the gene markers *TBR2* and *DLX2* or *DLX1*. *TBR2* (i.e. *EOMES*) reveals no expression in GE but strong expression cortical regions, while the reverse pattern is seen for *DLX2* and *DLX1* (**Fig. 18a, b, g, h**). Complementary expression patterns of *NKX2-1* and *ERBB4* are observed MGE and LGE, making it easy to distinguish them (**Fig. 18d, e**). The border between LGE and CGE is not sharp but CGE has much stronger expression of *CALB2* than LGE (**Fig. 13b**), and the reverse patterns occurs for *ERBB4* (CGE < LGE). The sulcus located between the striatum (GE) and the cortex, termed here striatal-cortical sulcus (SCS; indicated by arrows in **Fig. 18a-c, g-i**), does not appear to be a reliable landmark for the striatal-cortical border. As shown in **Figure 18**, the *NTRK2*-enriched striatal-cortical boundary (SCB, subpallial-pallial or pallial-subpallial boundary, see Puelles et al., 2000; Carney et al., 2009; **Fig. 17c, i**) is located at the striatal side at PCW 15 (**Fig. 18a-c**), while at PCW 21 it is at the cortical side (**Fig. 17g-i**). Also, the SCB is enriched with *CALB2, DLX2, ERBB4, PAX6* and *DLX1* (**Fig. 18b, e, f, h**).

**Fig. 18.**
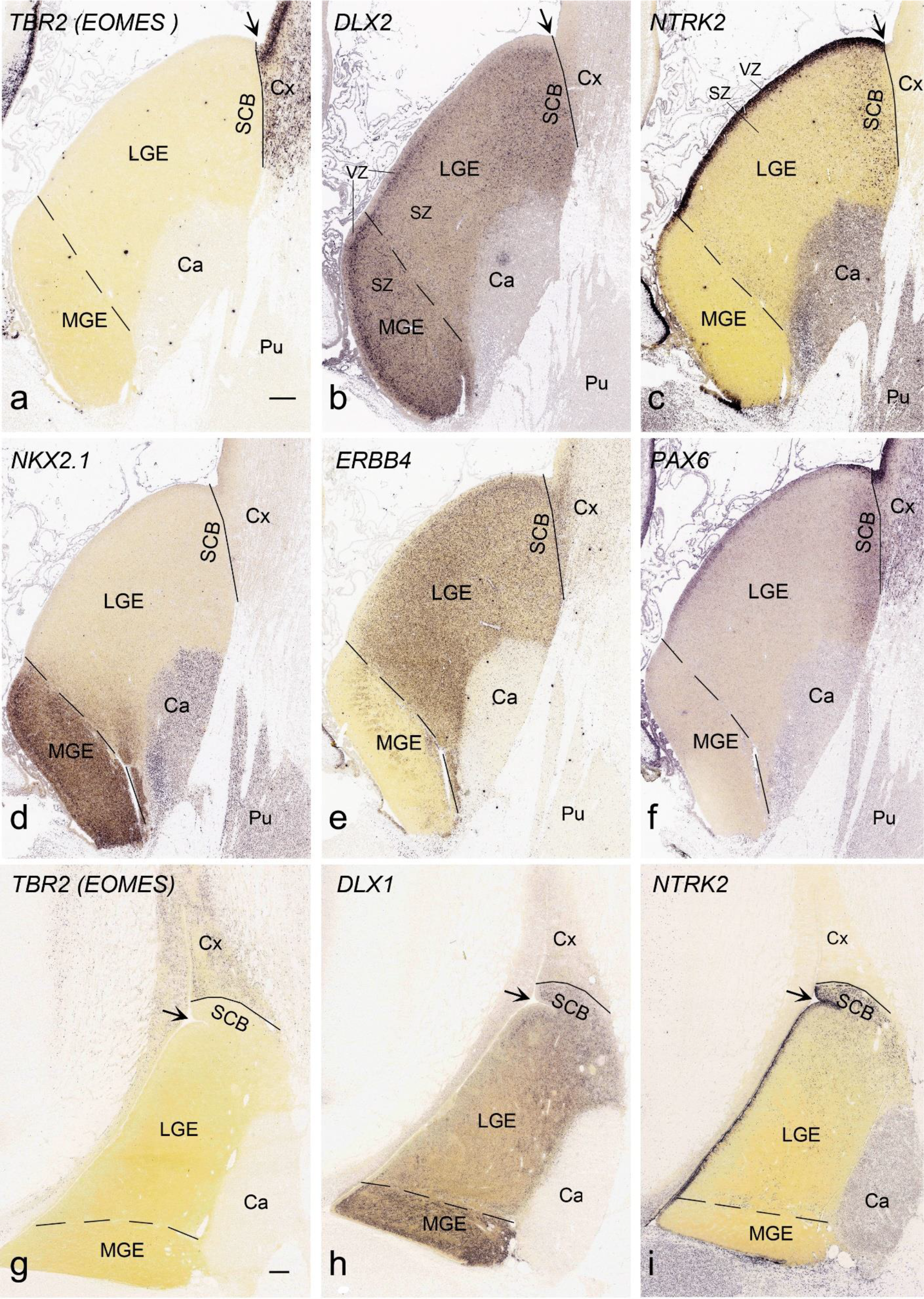
Identification of lateral and medial ganglion eminence (LGE and MGE) and striatal-cortical border (SCB). Solid and dashed lines indicate the striatal-cortical border and MGE-LGE border, respectively. VZ and SZ are ventricular and subventricular zones of the GE, respectively. (a-f). Gene expression in the GE and neocortex at PCW 15. *TBR2* (a) and *DLX2* (b) is expressed in the SZ/VZ of the cortex (Cx) and GE (LGE and MGE), respectively. *NTRK2* (c) and *PAX6* (f) are mostly expressed in the VZ of the LGE and SCB. *NKX2.1* (d) and *ERBB4* (e) are mainly expressed in SZ/VZ of MGE and LGE, respectively. Note *DLX2* and *ERBB4* expression also exists in SCB. (g-i). Gene expression in the GE and neocortex at PCW 21. Expression patterns of *TBR2* (g), *DLX1* (h), *NTRK2* (i), as well as *DLX2, NKX2.1, ERBB4* and *PAX6* (not shown) are similar to those at PCW 15. Surprisingly, the striatal-cortical sulcus (SCS, indicated by arrows) is not a reliable landmark for the striatal-cortical border. Thus, the SCB, as marked by rich NTRK2 expression, is located at the striatal side at PCW 15 (c), while at PCW 21 it is located at the cortical side (i). Scale bars: 400 µm in (a) for (a-f); 400 µm in (g) for (g-i).

In general, gene expression in the GE of the prenatal human brains is heterogenous with some gene expression in different zones within the GE. In MGE, for instance, distinct gene expression between its ventricular zone (VZ or inner part) and subventricular zone (SZ or outer part) and among different regions of the SZ is observed. Specifically, some genes are expressed in both VZ and SZ (e.g. *NKX2-1, NES* and *VIM;* e.g. **Fig. 18d**) with others exclusively or dominantly in SZ (e.g. *DCX, DLX5, DLX1* and *DLX2*; e.g. **Fig. 18b, h**) or VZ (e.g. *VIM*, *NTRK2* and *PAX6;* e.g. **Fig. 18c, f, i**). In LGE, heterogenous gene expression in its VZ and SZ is also observed for genes such as *NTRK2, FABP7* and *ERBB4*. Within the SZ of both LGE and MGE, region-dominant or complementary gene expression is visible, with stronger expression of *DCX, DLX1* and *DLX5* in the inner portion of the SZ than in the outer portion and a reverse expression pattern for *ERBB4, NES* and *VIM* (see **Appendix 2**).

### 3.10 Gene expression in olfactory bulb and anterior olfactory nucleus

At PCW 15, the olfactory bulb (OB) has obvious but immature lamination. From its outer to inner aspects, these layers include olfactory nerve (ON), glomerulus (GL), mitral cell (MC) and granular cell (GC), SZ and VZ. The SZ and VZ is part of the proliferative rostral migration system (RMS) (**Fig. 19a-p**). An incipient external plexus layer (ep) is also observed between GL and MC but the internal plexus layer is not well-defined. A number of genes show a strong expression in GL (*VIM, NTRK2, FADS2, SOX10* and *LMO4*), MC (*TBR2, GRIK2, LBX1, NTRK2, RELN, TBR1, LHX2, GAP43, DCX, DLX1, DLX5, NES, SYNGAP1, MECP2, SHANK3, FEZF2, CNTNAP2, GFAP, CDH4* and *LMO4*), GC (*DCX, FEZF2, LBX1, DLX1, SHANK3, CNTNAP2* and *LHX2*), SZ (*CALB2, DCX, PAX6* and *ZIC1*), and VZ (*VIM, TBR2, PAX6, FABP7, LHX2, FADS2, FOXG1, LBX1, ZIC1, ETV1* and *CDH4*). Interestingly, in addition to the OB lamination there exist patches within the OB (marked with # in **Fig. 19**) that are positive for *ENC1, GRIK2, LBX1, TBR1, LHX2. MECP2, CDH4, FEZF2, DCX, LMO4, PLXNA2, NKX2.1, FADS2,* and *GAP43*, and negative for *TBR2, RELN, CALB2, PAX6, ZIC1* and *DLX1* (**Fig. 19**).

**Fig. 19.**
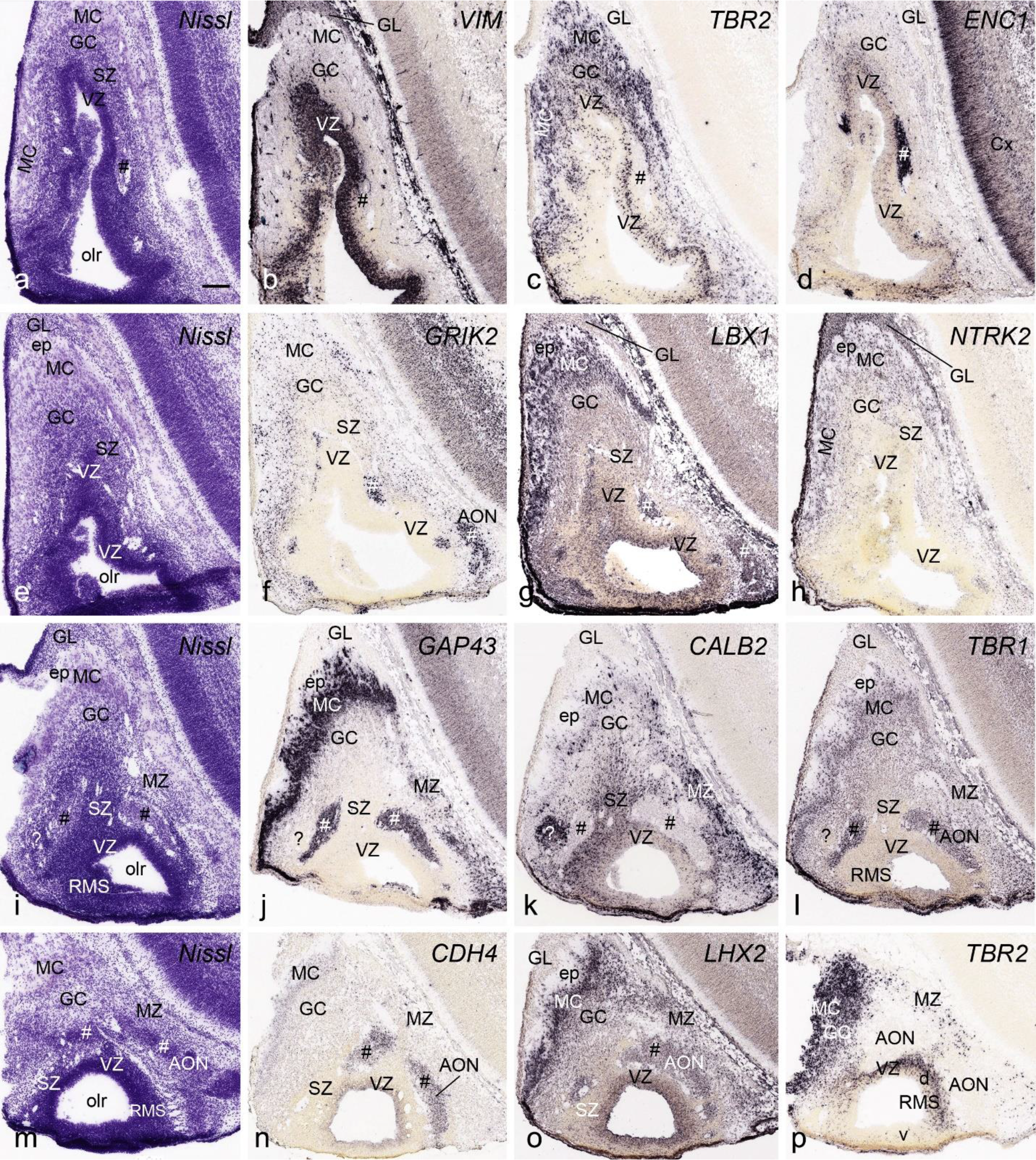
Cytoarchitecture and gene expression of olfactory bulb (OB) and anterior olfactory nucleus (AON) at PCW 15. The sections were shown from one rostral level (a-d), two intermediate levels (e-h and i-l) and one caudal level (m-p) with different stains indicated on each panel. OB, rostral migratory stream (RMS) and AON can be identified at PCW15. Some patches in OB and AON (identified by #) express *ENC1, GRIK2, LBX1, GAP43, TBR1, CDH4* and *LHX2* but not *CALB2*. Note that the dorsal but not ventral walls of the VZ of RMS strongly expresses TBR2 whereas the whole VZ of RMS contains strong expression of *VIM, LBX1* and *LHX2*. One patch (? in i-l) in AON appears to have a very different expression profile. MC, mitral cells; GC, granular cells; GL, glomerular layer; ep, external plexus layer; olr, olfactory recess. Bar: 200 µm in (a) for all panels.

The anterior olfactory nucleus (AON) is featured by strong expression of *ENC1, GRIK2, LBX1, GAP43, TBR1, CDH4, LHX2* and *MECP2* (**Fig. 19f, g, j, l, n, o**). The AON shows no expression of *PLXNA2, LMO4, TBR2, CALB2,* and *RELN* (e.g. **Fig. 19k, p**). Small patches are often seen in OB that have a pattern similar to that of the AON (e.g. **Fig. 19c-g**). The gene expression patterns observed in OB and AON at PCW 15 are comparable to those at PCW 21 (**Fig. 20**).

**Fig. 20.**
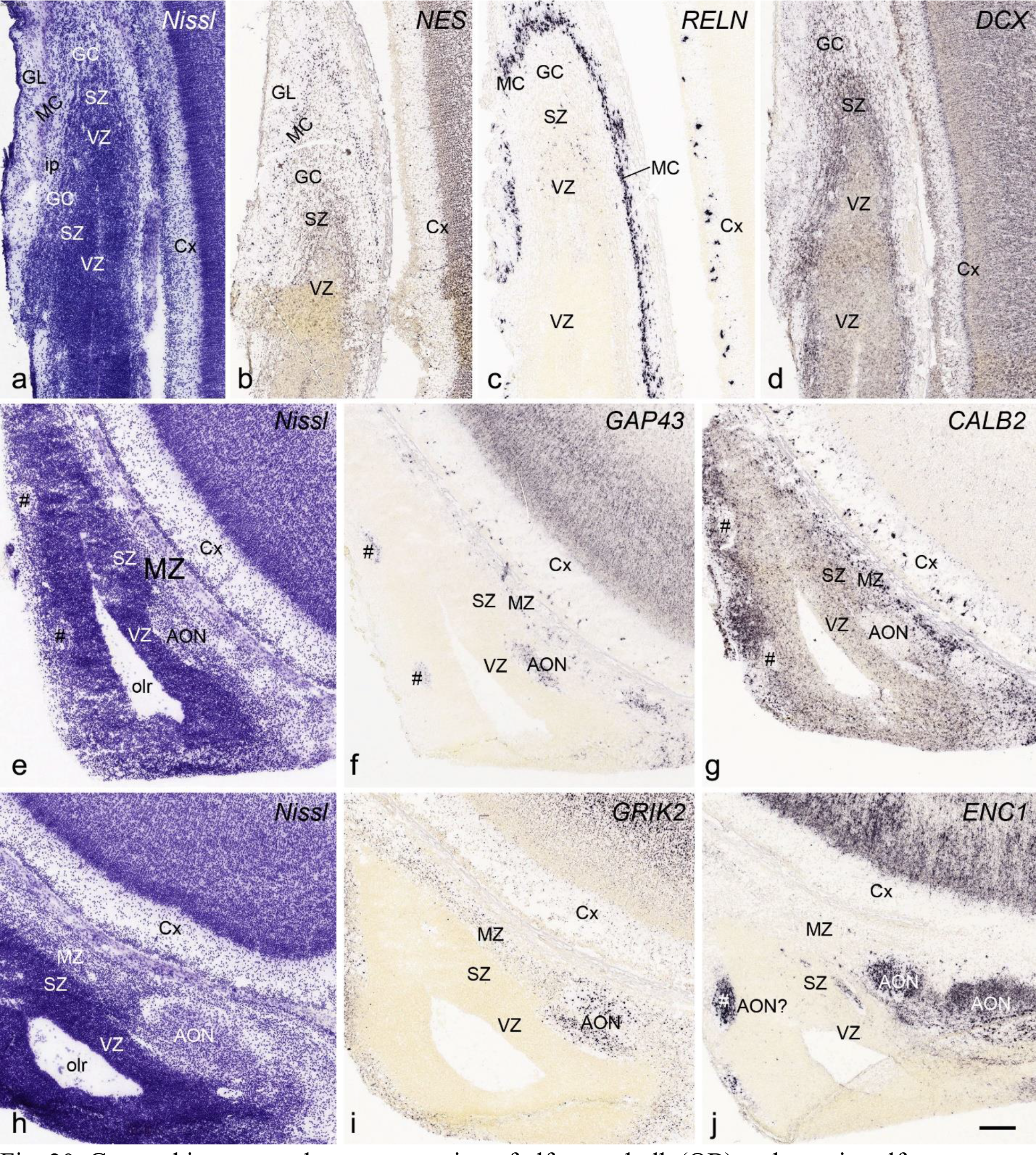
Cytoarchitecture and gene expression of olfactory bulb (OB) and anterior olfactory nucleus (AON) at PCW 21. The sections were shown from rostral (a-c), intermediate (d-g) and caudal (h-j) levels with different stains indicated on each panel. The layers of OB and internal plexus layer (ip) can be clearly identified at PCW 21. As at PCW15, AON strongly expresses *GAP43, GRIK2* and *ENC1* but negative for *CALB2*. Note the scattered patches (#s in e-g) which express marker genes for AON (e.g. *GAP43* and *ENC1*). Scale bar: 200 µm in (j) for all panels.

### 3.11 Brain-wide detailed anatomical and molecular atlases for prenatal human brains

Based on combined analysis of Nissl and AChE histology as well as laminar and regional gene expression patterns described above, anatomical boundaries of different cortical regions and subcortical nuclei can be accurately delineated on Nissl-stained sections. In this study, 46 and 81 selected Nissl-stained sequential coronal sections at PCW 15 and 21, respectively, were annotated in detail to generate two brain-wide anatomical atlases. The anatomical atlases for the brains at PCW 15 and 21 are presented in **Appendices 1 and 3**, respectively, with online links to high-resolution images. In these atlases, all cortical layers and many subcortical structures and their subdivisions are accurately demarcated based on cytoarchitecture and gene expression patterns. Furthermore, allocortex (HF and olfactory cortex), periallocortex (MTC and Iag) and their subdivisions are annotated. However, for neocortex only major cortical regions (e.g., frontal, parietal, temporal, occipital and insular cortices) could be roughly identified at PCW 15 as molecular makers did not reveal detailed regional patterns at this age although more detailed cortical segmentation could be generated for the brain at PCW 21. Finally, using the anatomical atlases as a guide, we have also annotated spatial expression of 37 and 5 genes from the brains at PCW 15 and 21, producing two brain-wide molecular atlases, which are presented in **Appendices 2** and **4**, with online links to high-resolution images. Although the expression of 38 genes from the brain at PCW 21 was not annotated the online links to their sequential high-resolution ISH images are presented in **Appendix 4**. Therefore, **s**patial mapping of these genes can be achieved by users using the detailed anatomical atlas (**Appendix 3**) as a guide, the ISH data and Nissl-stained sections used for the anatomical atlas being derived from the same brain hemisphere.

### 3.12 Highly interactive digital atlases for web users

The prenatal brain atlases presented in this study were used to make interactive digital resources (**Fig. 21**), and are publicly accessible through the Allen Institute web portal: www.brain-map.org or directly at the BrainSpan project portal: www.brainspan.org/static/atlas. The atlases may be of interest to diverse groups, including students and educators as resources of detailed anatomy of the prenatal human brains. For basic anatomy, the location, shape and relationship of general structures such as forebrain, midbrain and hindbrain, as well as cerebral cortex, thalamus, hypothalamus, amygdala, hippocampus, cerebellum and brainstem can be easily elucidated simply by using the ontology tree (**Fig. 21, left column**). For medical students and neuroscience professionals, deeper systematic learning and teaching of human brain development can be conveniently achieved via the menu and different tools (**Fig. 21, inset, thumbnails and right column**). In particular, one can search or choose any specific brain region and learn its topographic location, subdivisions, cyto- and chemo-architectures. One can also choose specific A-P levels and study which structures occurs on these planes. As needs dictate, structures can be recolorized prior to printing plates using the menu (**Fig. 21, inset**). The ontology itself serves as an essentially complete index of brain structures and their relationships (**Fig. 21** left column; **Table 1**).

**Fig. 21.**
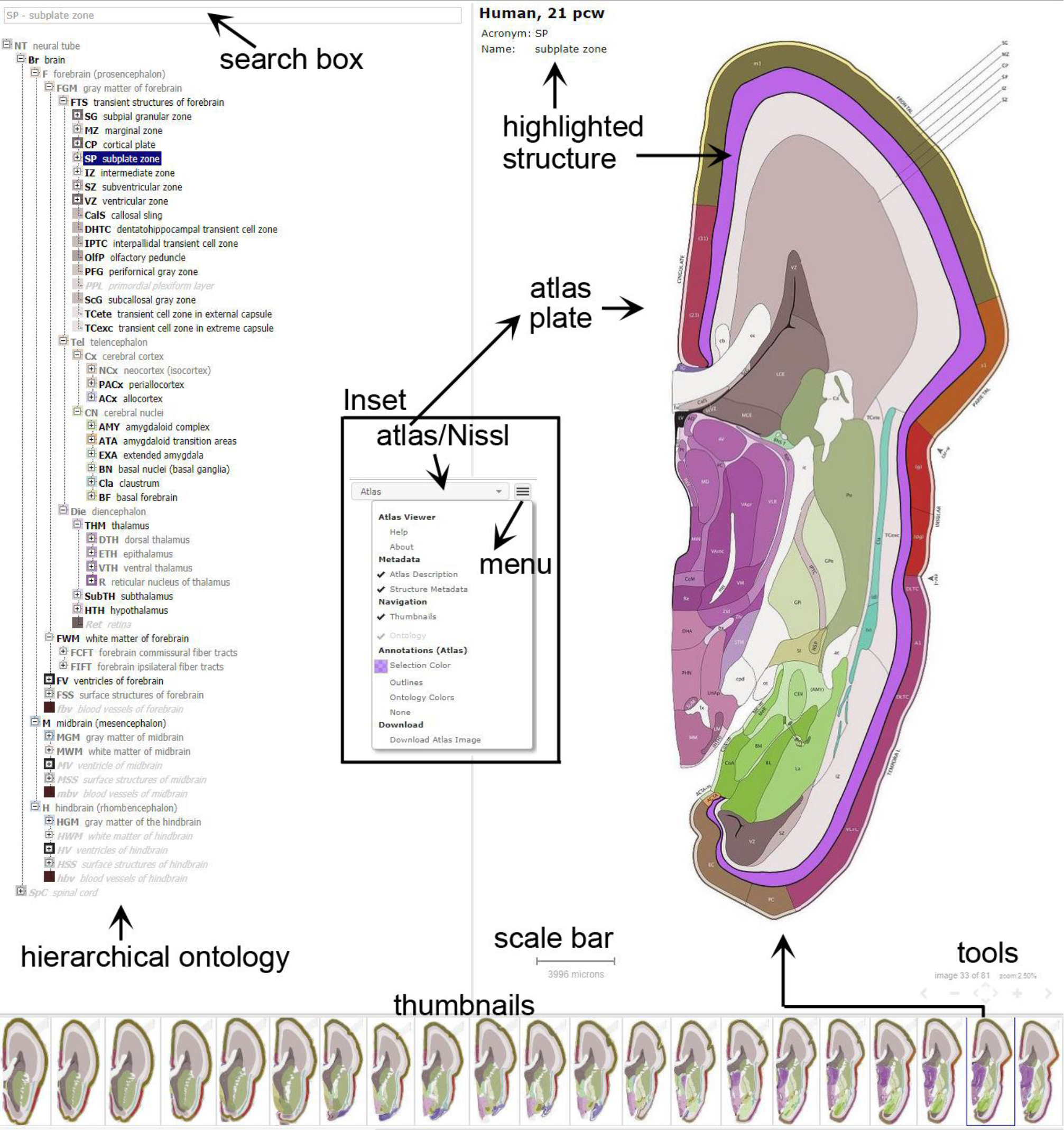
Interactive web-based digital atlases for prenatal human brains at PCW 15 and 21 (see the links below). Tools are provided to explore, search, adjust and download the atlas plates and related structures. These tools include a hierarchical ontology browser and search box (left column), thumbnails (bottom), segmented atlas plates (right column), and ability to toggle between Nissl and annotated plates.

## DISCUSSION

Anatomical atlases are essential resources as for the research community, providing a detailed mapping and synthesis of knowledge about brain structure and function. Although detailed modern high-resolution human brain atlases are available for adult brains (Ding et al., 2006; Mai et al., 2016), similar atlases for prenatal human brain have not been produced. To our knowledge, only one series of anatomical prenatal human brain atlases is available, generated on limited Nissl-stained sections from different prenatal ages (Bayer and Altman, 2003; 2005; 2006). In addition, anatomical delineation in these developmental brain atlases was based only on cytoarchitecture (Nissl staining), and annotation was not complete enough to create a brain-wide hierarchical structural ontology. In the present study, we have created detailed anatomical atlases for prenatal human brain at PCW 15 and 21, aiming to advance the state of the field through dense whole-brain sampling, high information content histological and gene expression analysis, developmental ontology creation, and generation of web-based interactive tools. This reference atlas was used to guide a large-scale microarray-based transcriptomic project, available as a complementary developmental gene expression resource (Miller et al., 2014; https://www.brainspan.org/lcm/search/index.html; see Table 1 for the regions with transcriptomic data). Thus, these atlases should be valuable tools to guide newer efforts to map cell types and developing circuitry in the developing human brain, both as anatomical and gene expression resources.

A major design principle of the atlas was to use highly informative histological and gene expression datasets from the same brain specimen to guide anatomical demarcation across the entire brain at PCW 15 and 21. As demonstrated above, the combined analysis of Nissl-based cytoarchitecture, AChE staining features and spatial expression patterns of 43 genes by ISH allowed an accurate delineation of brain structures. This information was used to annotate Nissl-stained sections, but also to annotate the gene expression images and illustrate the high utility of individual genes as markers of developing brain structures. Together, these atlases provide a reference for mid-gestational human brain development and can be used to annotate other timepoints in this period. For example, we used the anatomical atlas for the brain at PCW 15 (**Appendix 1**) to guide a dense transcriptomic atlas of the PCW 16 brain, which was effective due to its similar cortical lamination and subcortical structures (see Miller et al., 2014). The anatomical atlas for the brain at PCW 21 (**Appendix 3**) was similarly created and could be applied to the human brains at ages close to PCW 21 (e.g., PCW 19-24). With guidance of these anatomical atlases, spatial expression of 37 and 5 genes was annotated to generate two brain-wide molecular atlases, which are presented in **Appendices 2** and **4,** respectively.

Several interesting developmental features of prenatal human brain emerged from this analysis. First, we found that the prenatal HF and EC lack the outer subventricular zone (SZo) that is prominent in developing human neocortex and thought to drive the differential expansion of supragranular layers in primate evolution. The SZo is the major site of supragranular neuron production in the macaque monkey neocortex (Lukaszewicz et al., 2005), and is also present (albeit smaller) in rodents, where common gene expression in SZo and supragranular neurons suggests they are generated from SZo (Nieto et al., 2004; Tarabykin et al., 2001; Zimmer et al., 2004). The thicker SZo in human compared to mouse is also consistent with single-cell transcriptomic findings of increased number, diversity and phenotypic specialization of supragranular neurons (Ortega et al., 2018; Berg et al., 2021). Histologically, the HF and EC obviously lack equivalent supragranular neurons in layers 2 and 3 of the neocortex (Bakken et al. 2016; Ding & van Hoesen, 2015). While layers 2 and 3 exist in the EC the neurons in these two layers are not equivalent to those in the neocortex. For example, layer 2 neurons in the neocortex are very small round and ovoid neurons, while those in the EC are usually very large stellate or pyramidal neurons (Braak & Braak, 1991; Ding & van Hoesen, 2010). Gene expression and connectivity patterns of layers 2 and 3 in the EC are also different from the neocortex (e.g. Bakken et al., 2016; Ding et al., 2020; Yao et al. 2021). Pathologically, layer 2 neurons in the EC are among the earliest neurons with tau lesions in aging and Alzheimer’s disease (AD) populations (Arnold et al. 1991; Braak & Braak, 1991; Ding & van Hoesen, 2010). However, this is not true for layers 2 and 3 neurons in the neocortex, which are affected at later stages of AD (Braak & Braak, 1991). Therefore, our findings suggest that layers 2 and 3 neurons in the PrS, PaS and EC are not likely produced from SZo and instead may directly originate from the SZi. As a previous study suggested, the hippocampal neurons are generated in the VZ while MTC neurons are generated in the VZ and SZ (SZi) with the latter generating the superficial neurons located in lamina principalis externa (Nowakowski & Rakic, 1981).

We found evidence for compartmentalization of the ganglionic eminence (GE) in developing human cortex, similar to multiple progenitor domains that have been reported in mouse GE (Flames et al., 2007). In addition to the three well-known subdivisions of the GE (MGE, LGE, CGE), each subdivision can be further divided into VZ and SZ parts that continue at the SCB with cortical VZ and SZ, respectively. The VZ and SZ of the GE displayed differential gene expression within each GE subdivision, and cross over between GE subdivisions (e.g., **Fig. 18**). Moreover, complex gene expression patterns within the VZ and SZ of specific GE subdivisions in human were also clearly evident. For example, *NTRK2* is predominantly expressed in the VZ of CGE and LGE; furthermore, within the LGE, *NTRK2* expression in the VZ is generally stronger in lateral LGE (and SCB) than in medial LGE (see **Appendix 2**). Gene expression in the SZ of LGE is also not homogeneous. For instance, *FABP7* expression in the SZ of the LGE is stronger in lateral than medial parts (see **Appendix 2**), while *NKX2.1* expression is seen in the most medial but not lateral parts of the SZ in addition to strong expression in MGE. Interestingly, *ERBB4* expression displays both A-P and M-L difference in the SZ of the LGE. Specifically, *ERBB4* is much more strongly expressed in the SZ of the latero-caudal part of the LGE (including SCB) compared to the medio-rostral part (see **Appendix 2**). Finally, *SST, NPY, VIM, FABP7* and *ERBB4* expression is also compartmentalized in MGE (see **Appendix 2**). These observations likely represent a combination of parcellation of progenitor zones and developmental gradients across the GE.

*ERBB4* expression in the GE is significantly different between human and rodents. Previous studies in rat and mouse reported that *ERBB4* is mostly expressed in the SZ of MGE (i.e. not in LGE and CGE) (Yau et al., 2003; Fox & Kornblum, 2005). Rodent *ERBB4* is mainly expressed in MGE-derived GABAergic interneurons migrating to the cerebral cortex, and this expression was reported to be important in this tangential migration (Li et al., 2012; Rakic et al. 2015; Villar-Cerviño et al., 2015). In contrast, we found in human that *ERBB4* is strikingly enriched in LGE rather than MGE. This raises the possibility that human LGE generates cortical interneurons that express *ERBB4* and migrate tangentially through the SZ and VZ (see **Appendix 2**). Alternatively, this may simply reflect a species difference in *ERBB4* expression such that OB-bound interneurons from the LGE now express *ERBB4*, or that *ERBB4*-expressing interneurons are mostly generated in MGE as in rodents, but they are immediately channeled to LGE in human. In any event, these findings in human suggest that there may be significant differences in MGE and LGE across species, and that at a minimum *ERBB4* expression and function are not conserved across species.

Another interesting observation was that markers of both cortical and striatal progenitors extend into the rostral migratory stream. In mouse, the RMS is a pathway mostly consisting of interneurons migrating from LGE and SCB to OB (e.g., Kohwi et al., 2005; Bandler et al., 2017). Glutamatergic neurons also migrate in the RMS, but the source of these neurons has not been clear. In mice, these excitatory neurons derive from *NGN2*-expressing progenitors (Winpenny et al. 2011), and excitatory lineage markers *NGN2*, *PAX6* and *TBR2* are all expressed in the dorsal-medial wall of the RMS (RMS-dm) but also in the SCB region. Here we found that *TBR2* (i.e., *EOMES;* Englund et al, 2005; Hevner, 2019) is strongly expressed in the developing cortical VZ and SZ and in the RMS-dm, but not the SCB region. These findings suggest that the *TBR2*-expressing VZ and SZ of the neocortex extends rostrally into the OB via the RMS-dm, and that excitatory neurons in OB likely originate from the ventral cortical wall rather than from the subpallial-pallial region or SCB region, both during development (this study) and into adulthood (Brill et al., 2009).

Single cell genomic profiling has become a powerful tool to define cell types (Hodge et al., 2019; BICCN, 2020; Yao et al., 2021) and developmental trajectories, and has begun to be applied in prenatal human brain development (Nowakowski et al., 2017; Fan et al., 2020; Eze et al., 2021). Whole-brain anatomical and molecular atlases such as those presented here are important resources to help guide these new cell census efforts and other high-throughput anatomical and connectional efforts in the future. The joint analysis of histology and molecular parcellation creates a needed framework for establishing the cellular and spatial basis of brain development and circuit formation, and a structured ontological framework for defining brain structures across development to adulthood.

## ACKNOWLEDGMENTS

We thank the Allen Institute founders, P.G. Allen and J. Allen, for their vision, encouragement, and support. We are also grateful for the technical support of the staff members in the Allen Institute who are not part of the authorship of this paper. The project described was supported by award numbers RC2MH089921 to ESL and MJH from the National Institute of Mental Health. Its contents are solely the responsibility of the authors and do not necessarily represent the official views of the National Institutes of Health and the National Institute of Mental Health.

## AUTHOR CONTRIBUTION

SLD and ESL designed the atlas project; SLD generated the anatomical ontology, analyzed histological and gene expression data and delineated all anatomical boundaries and spatial gene expression. JJR, PL and BACF performed atlas cartography; KAS led the ISH experiment; IAG, SLD and ND contributed to specimen processing; TAD, NHK, FL, AS and LN performed web visualization and application; KB, RAD, AE, KG, TAL, JN, JP, RR, MS and NVS provided technical support; YW, SLD, JJR, PL and BACF prepared appendix figures; SLD prepared the figures for the main text; SMS managed the project; AB managed and led the histological pipeline and team; JWP, JGH, ARJ, MJH, PRH, AB and ESL provided overall supervision. ESL and MJH procured the NIH grant; SLD and ESL wrote the manuscript; all authors had full access to all the data and read the manuscript.

## DATA AVAILABILITY STATEMENT

The data that support the findings and atlas creation of this study are available online (www.brain-map.org or www.brainspan.org), and available from the corresponding authors upon reasonable request.

## Appendix 1 Anatomical atlas plates for the prenatal human brain at PCW 15

Forty-six plates with matching histological (Nissl- or AChE-stained) sections are displayed in anterior-to-posterior (A-P) order. The A-P positions of 46 atlas plates (levels 1-46) and the locations of the two slabs as well as the codes for A-P levels, stains, slabs and the section number in each slab are indicated in the Inset on top of the first page of the atlas plates. For example, Level 1 (01–015) represents the 15th section of the slab 1, which is atlas plate 1 (i.e., Level 1), and AChE-01-054 represents the 54th section of the slab 1, which is AChE-stained and adjacent to plate 2 [Level 2 (01-053)]. The parcellation and subdivisions of different brain regions as well as the parent–daughter relationship and abbreviation of each structure are detailed in **Table 1** (for transient structures) and in our adult human brain atlas (Ding et al. 2016; for permanent structures). Note that all annotated brain structures and their abbreviation in this atlas can be searched in the interactive window of the online version of this atlas (www.brainspan.org or http://brainspan.org/static/atlas).

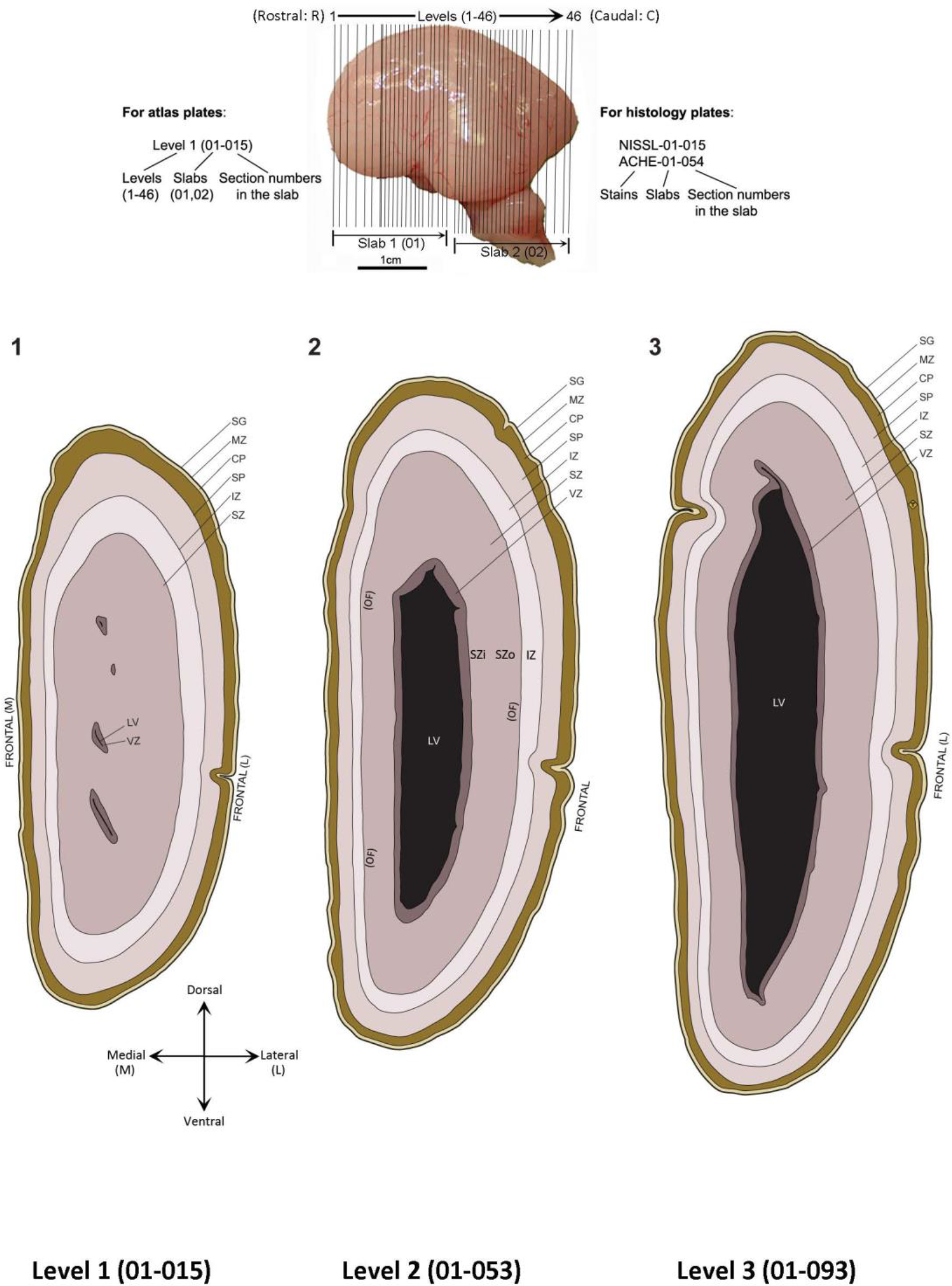
Appendix 1 (showing first two atlas plates as examples):

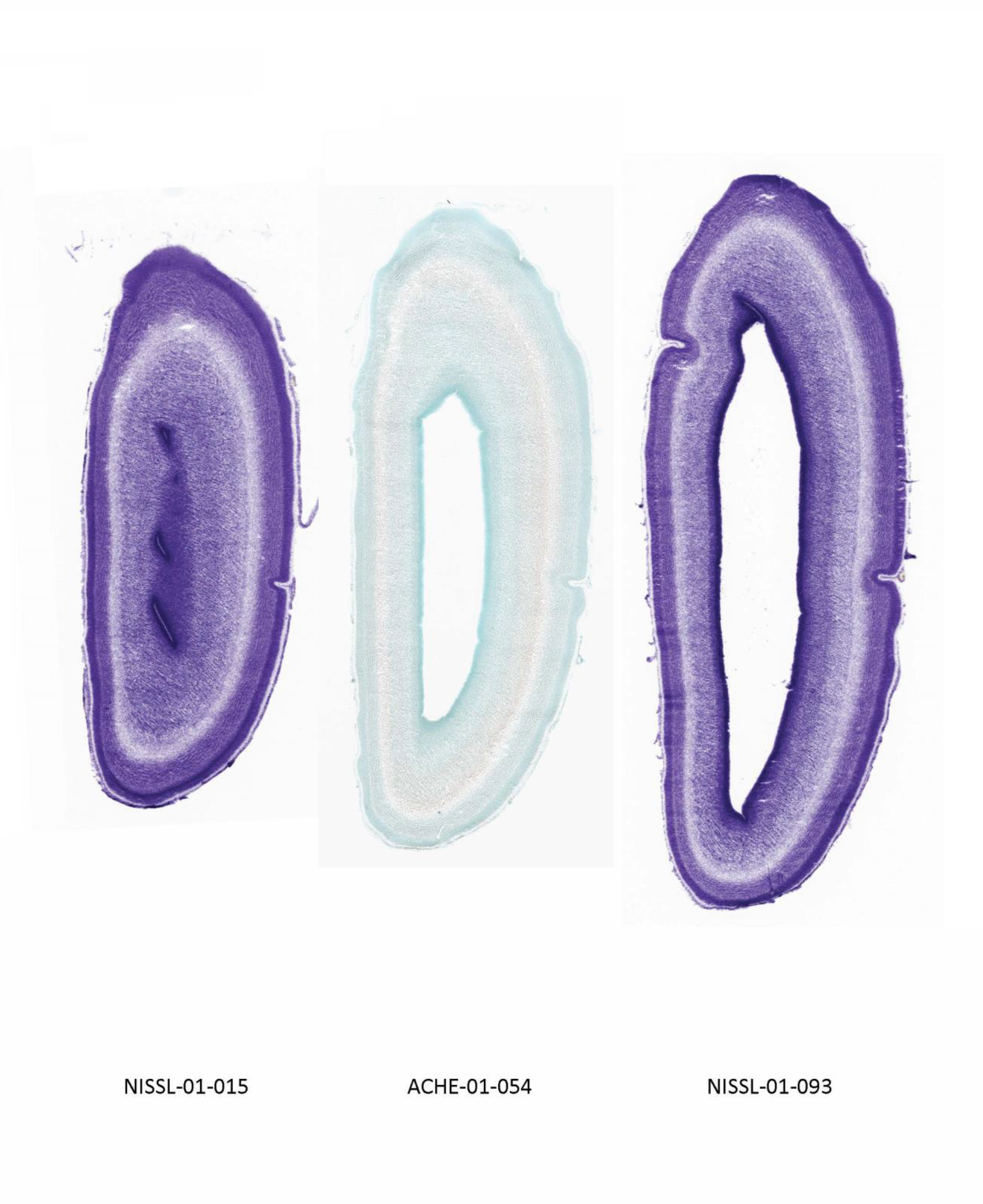

## Appendix 2: Spatial expression of 37 genes in the brain at PCW 15

Anatomical localizations of the gene expression are based on the anatomical brain atlas (**Appendix 1**) which is derived from the same brain hemisphere. Note that some artifacts like the blank (white) round or oval regions usually caused by air bulbs during ISH staining are indicated by (#). The expression of the other 6 genes was not annotated mainly because of their faint or sparse expression but the sequential ISH images for **these** genes are linked online after the 37 genes in this appendix.

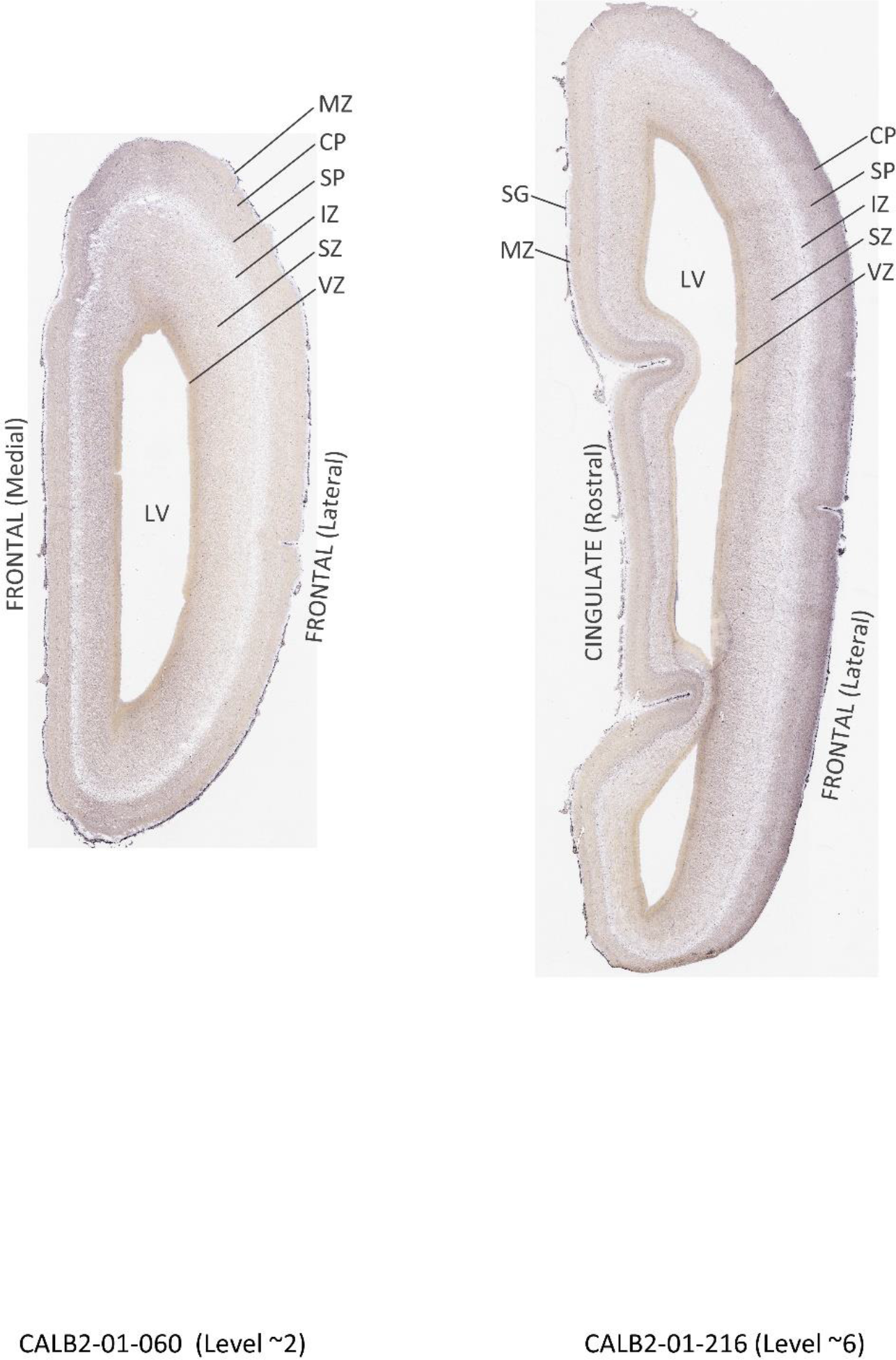
Appendix 2 [showing one gene (CALB2) as an example]:

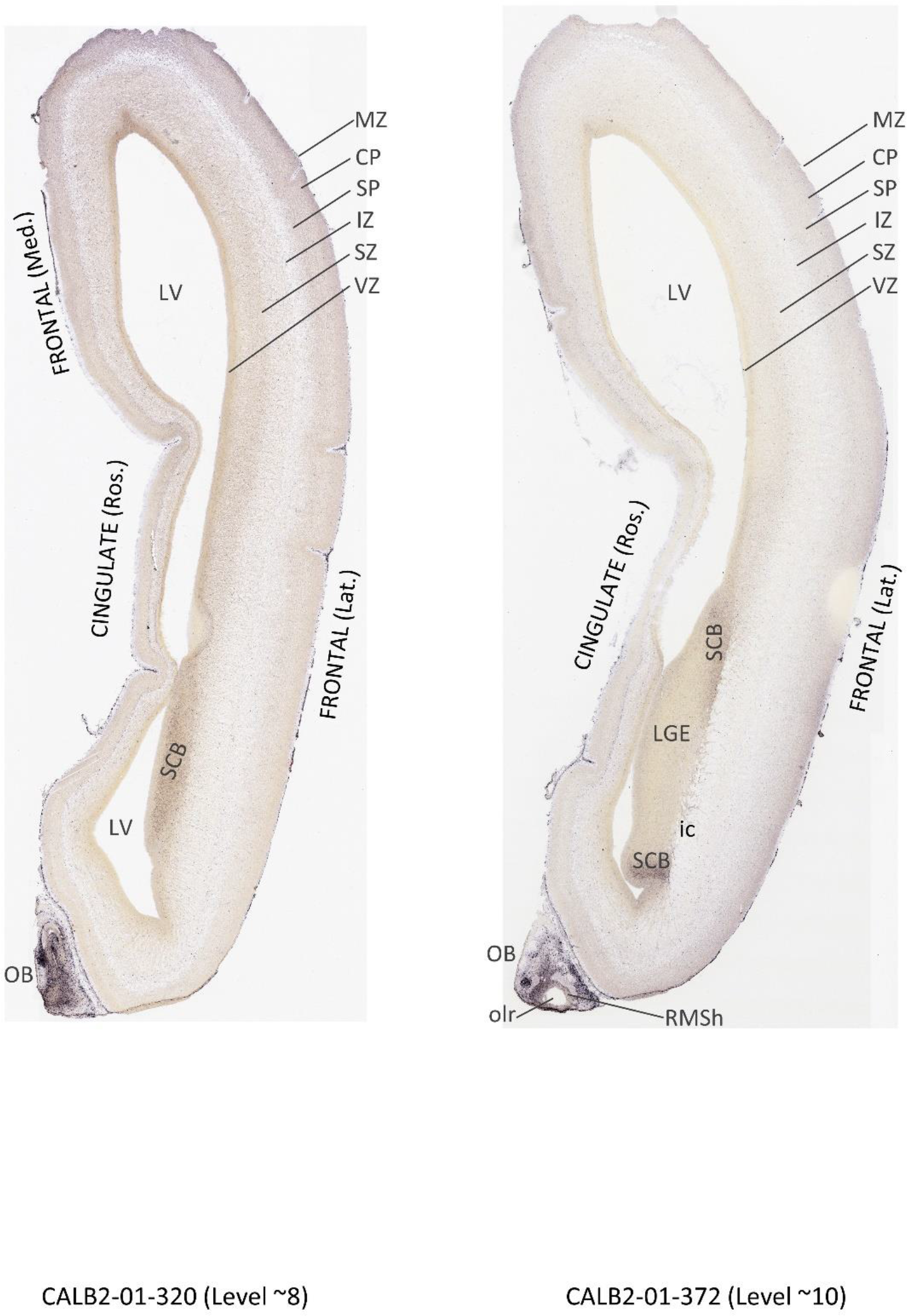

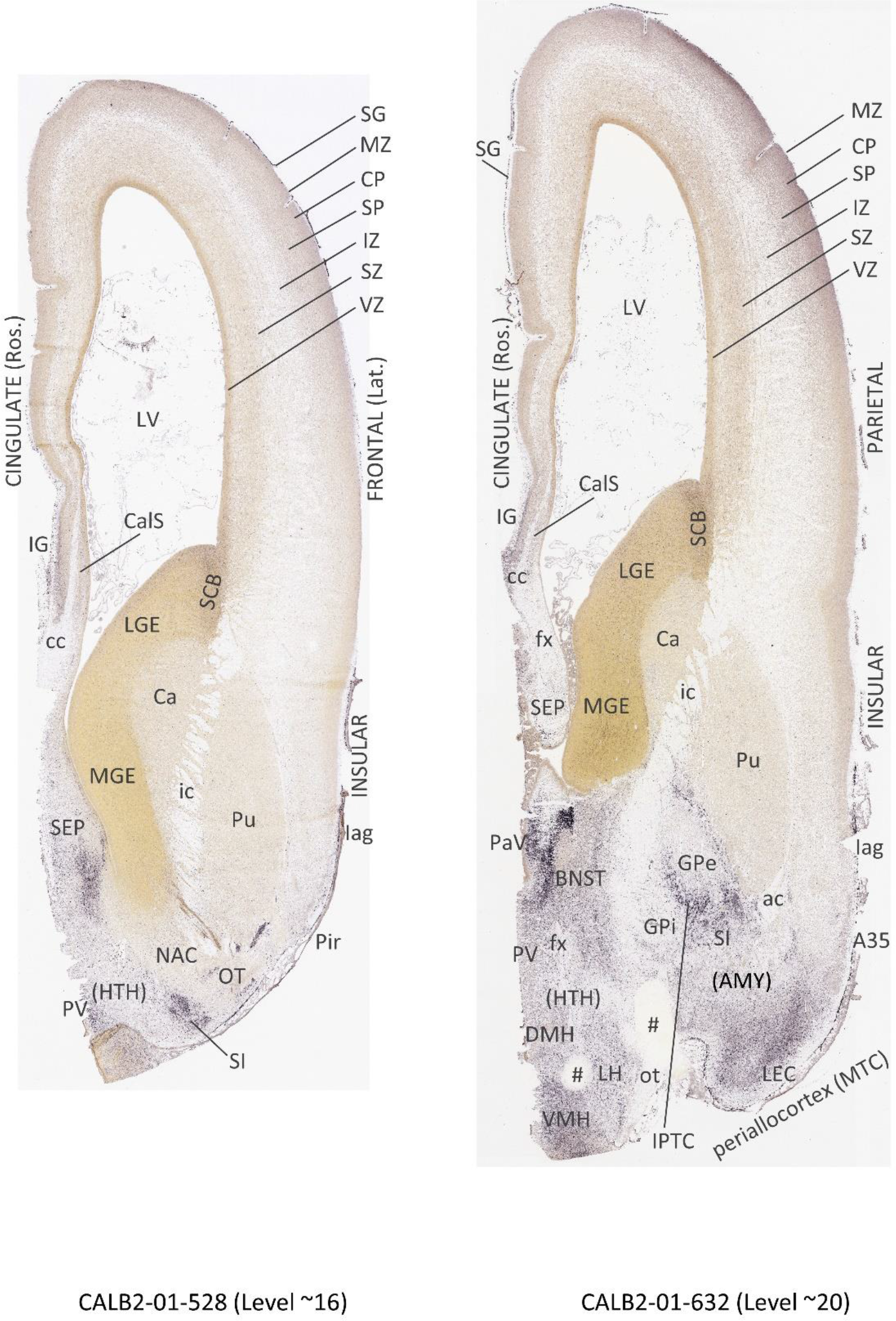

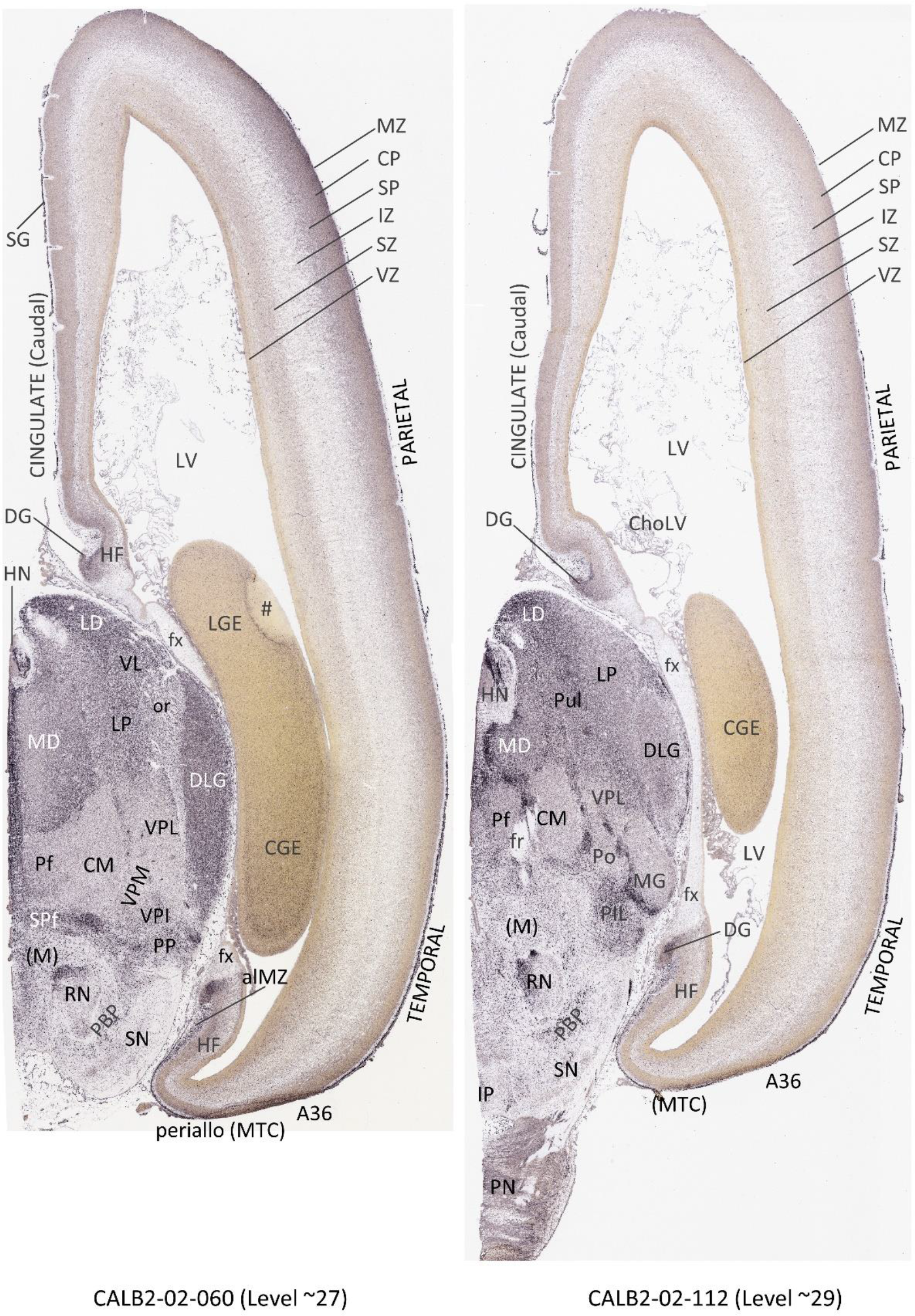

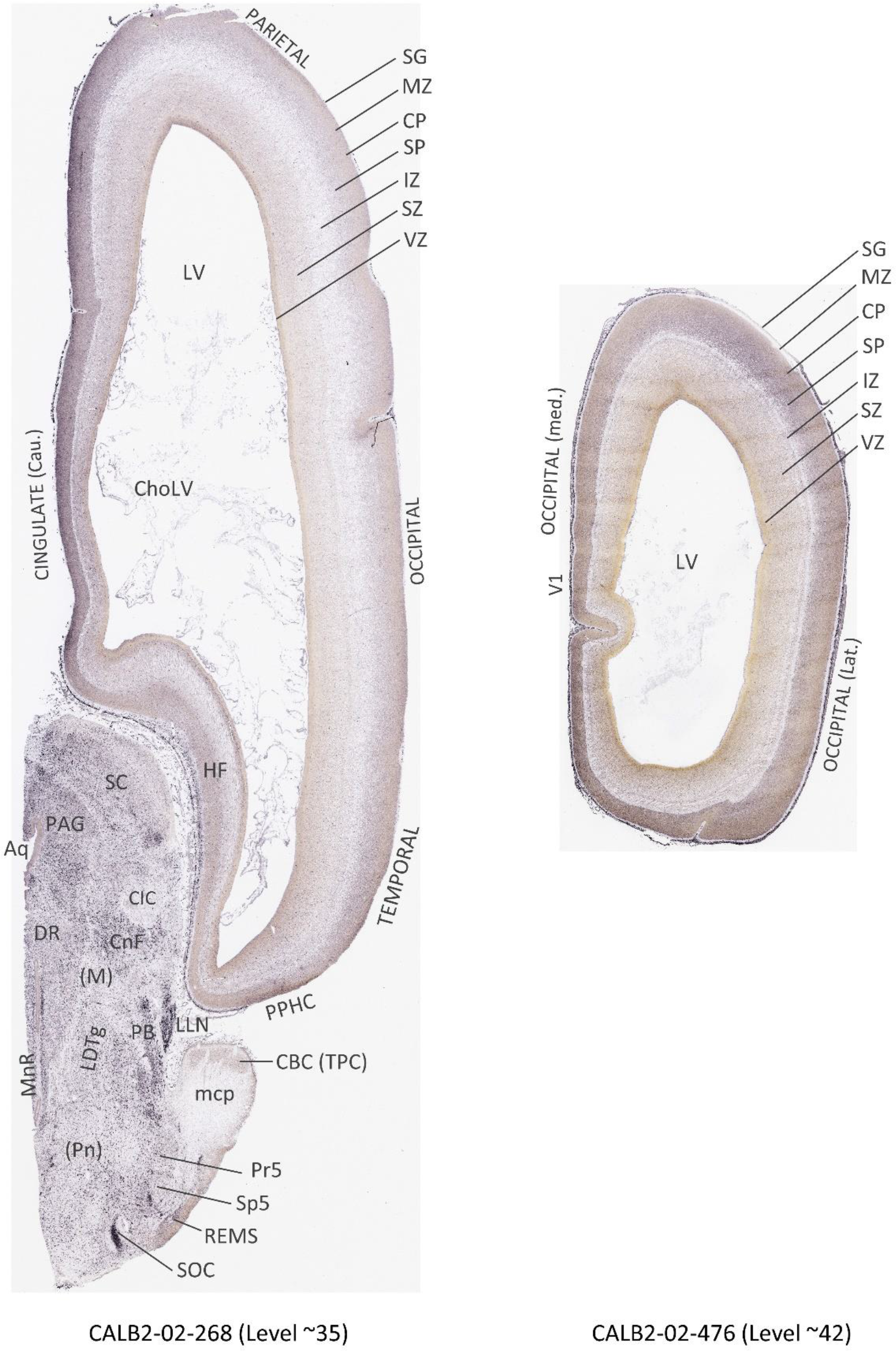

## Appendix 3: Anatomical atlas plates for the prenatal human brain at PCW 21

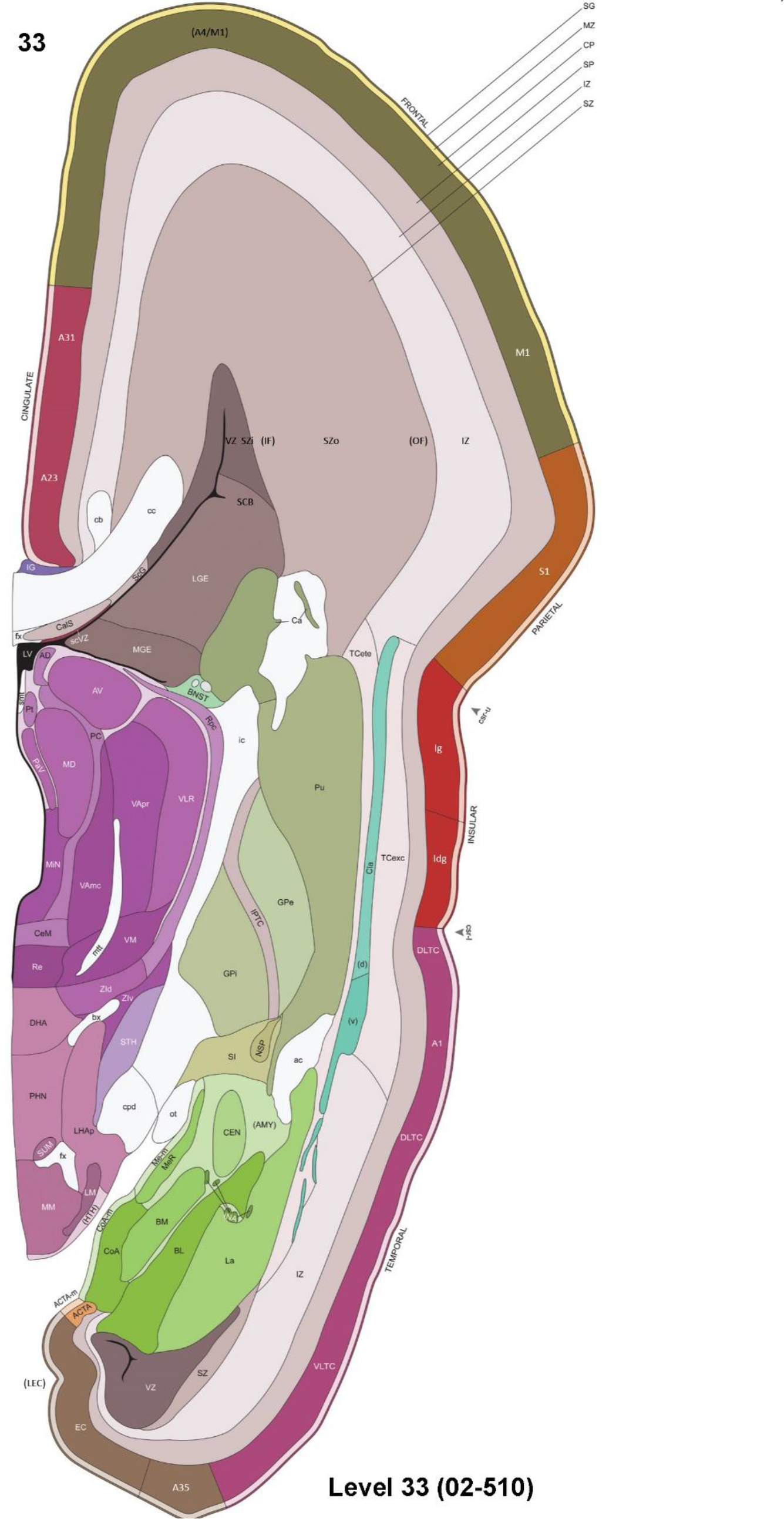
Appendix 3 (showing two atlas plates as examples):

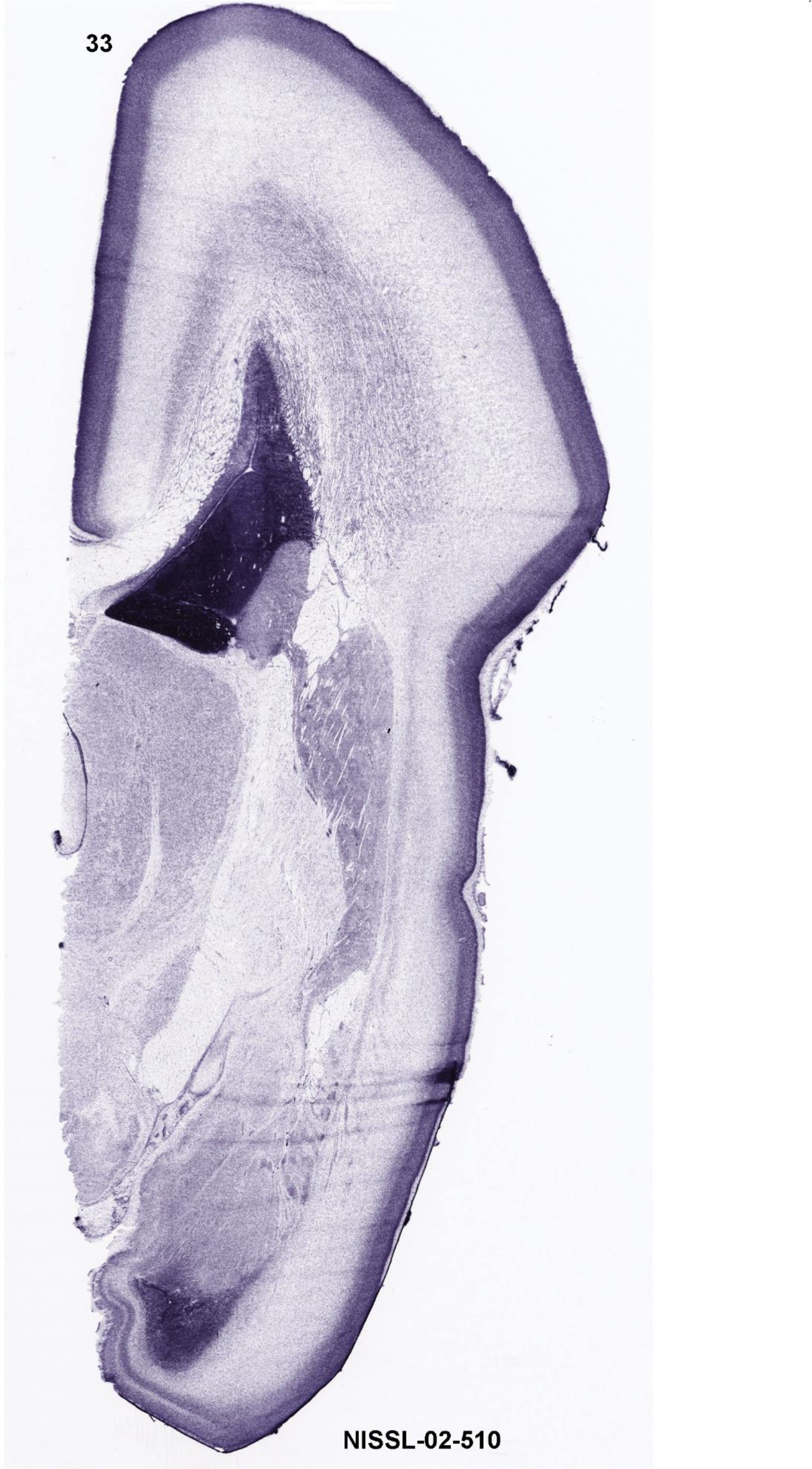

Eighty-one plates with matching histological (Nissl-stained) sections are displayed in anterior-to-posterior (A-P) order. The A-P positions of 81 atlas plates (levels 1–81) and the locations of the four slabs and their the starting and ending plate levels in each slab are indicated in the Inset of this appendix (below). For each atlas plate, its A-P levels, slab and section numbers are labeled in this style: Level number (slab number-sections number in the slabs). For example, Level 1 (01–026) represents the 26th section from slab 1, which is atlas plate 1 (i.e. Level 1). The matching Nissl-stained sections are labeled in this way: NISSL-slab number-section number in the slabs. For example, NISSL-01–026 represents the 26th section from slab 1, on which the annotation of atlas plate 1 (Level 1) was performed. In addition, one series of sequential AChE-stained sections from the same brain hemisphere is also available and linked online here (link). The parcellation and subdivisions of different brain regions as well as the parent–daughter relationship and abbreviation of each structure are detailed in **Table 1** (for transient structures) and in our adult human brain atlas (Ding et al. 2016; for permanent structures). Note that all annotated brain structures and their abbreviation in this atlas can be searched in the interactive window of the online version of this atlas (www.brainspan.org or http://brainspan.org/static/atlas).

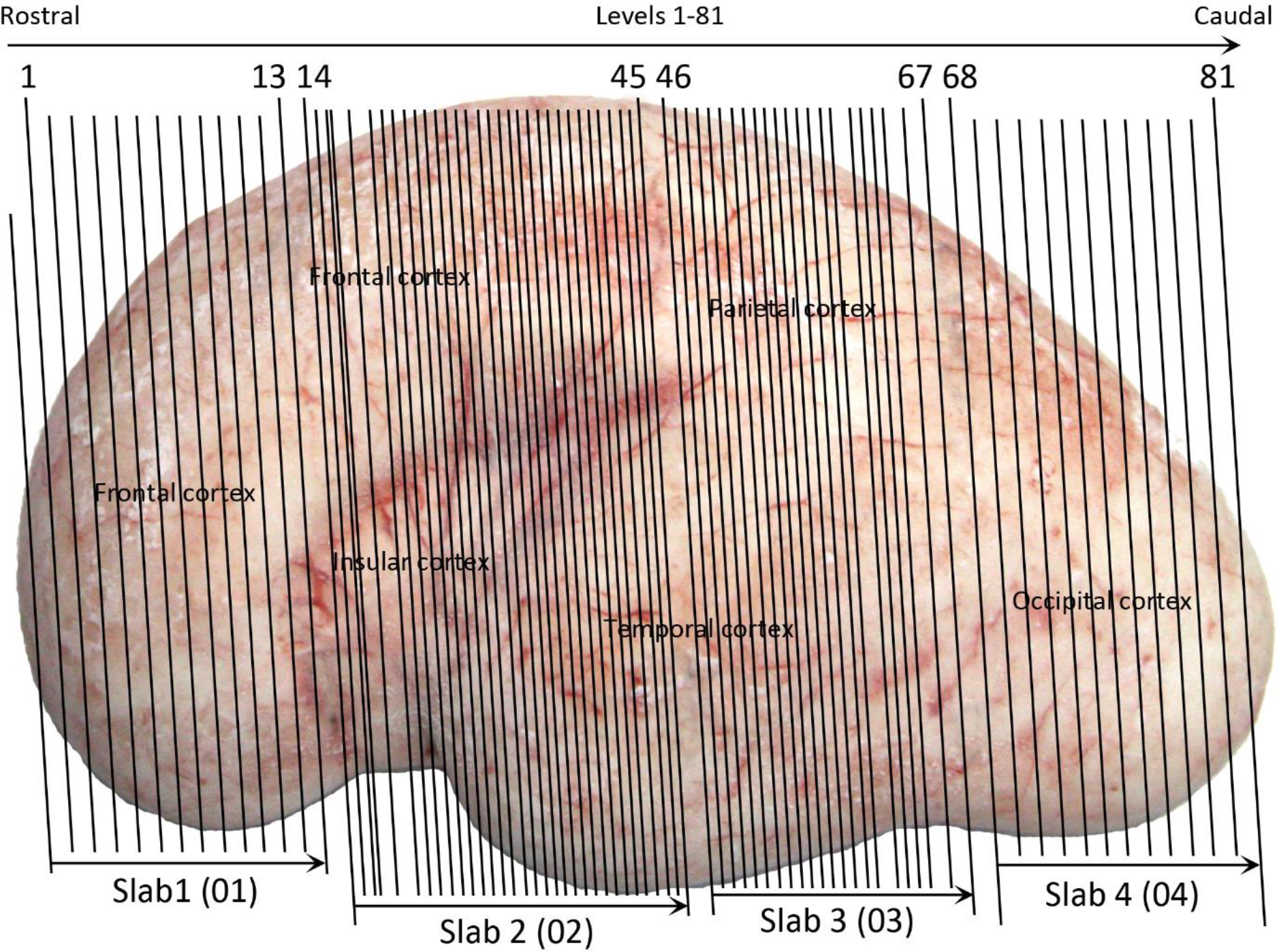

## Appendix 4: Spatial expression of 5 representative genes in the brain at PCW 21

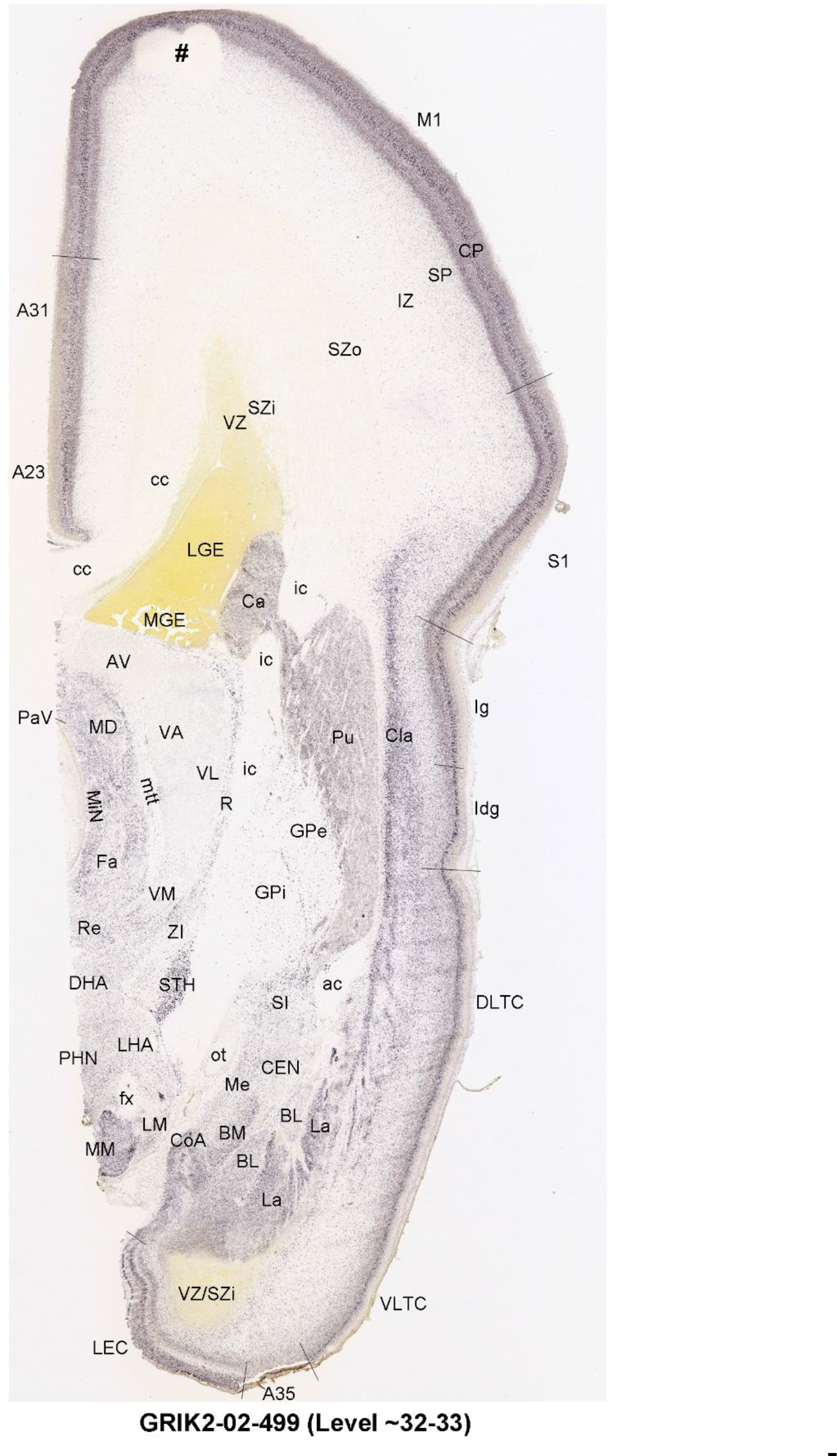
Appendix 4 [showing one GRIK2-ISH plate as an example]:

Anatomical localizations of the gene expression are based on the anatomical brain atlas (**Appendix 3**) which is derived from the same brain hemisphere. Note that some artifacts like the blank (white) round or oval regions usually caused by air bulbs during ISH staining are indicated by (#). The spatial expression of the other 38 genes was not annotated because of limited length of this paper but the sequential ISH images of these genes are linked online in this appendix. The spatial locations of their expression can be easily determined using our anatomical brain atlas (**Appendix 3**) as a guide, as demonstrated for the 5 genes displayed here.

